# Electrical impulse characterization along actin filaments in pathological conditions

**DOI:** 10.1101/2021.09.03.458927

**Authors:** Christian Hunley, Md Mohsin, Marcelo Marucho

**Author notes:** Email address:* (Marcelo Marucho).

## Abstract

We present an interactive Mathematica notebook that characterizes the electrical impulses along actin filaments in both muscle and non-muscle cells for a wide range of physiological and pathological conditions. The program is based on a multi-scale (atomic → monomer → filament) approach capable of accounting for the atomistic details of a protein molecular structure, its biological environment, and their impact on the travel distance, velocity, and attenuation of monovalent ionic wave packets propagating along microfilaments. The interactive component allows investigators to conduct original research by choosing the experimental conditions (intracellular Vs in vitro), nucleotide state (ATP Vs ADP), actin isoform (alpha, gamma, beta, and muscle or non-muscle cell), as well as, a conformation model that covers a variety of mutants and wild-type (the control) actin filament. The simplicity of the theoretical formulation and the high performance of the Mathematica software enable the analysis of multiple conditions without computational restrictions. These studies may provide an unprecedented molecular understanding of why and how age, inheritance, and disease conditions induce dysfunctions in the biophysical mechanisms underlying the propagation of electrical signals along actin filaments.

## 1. Introduction

Actin filaments are a major component of the cytoskeleton, and essential for various biological activities in eukaryotic cellular processes such as directional growth, shape, division, plasticity, and migration [1, 2, 3, 4, 5]. Recent research on the electrical properties of actin under in vitro conditions revealed an unprecedented role for the microfilaments as bionanowires capable of propagating ionic information [6]. This finding may provide new insights on many electrical processes taking place within smooth muscle, skeletal, and cardiac cells near the membrane or in the nucleus. Furthermore, an additional information delivery system in neuronal cells might be beneficial for intracellular communication in dendrites, the soma, axon, and axon terminal [7]. Indeed, further clarification on the electrical conductivity and ionic transmission properties of actin filaments is vital for a complete understanding of their contributions to cellular functions and information transfer inside cells.

The use of conventional computational tools and approaches to study the interplay between the polyelectrolyte properties of polymer chains and their biological environment break down for cytoskeleton filaments, because they are limited by their approximations and computational cost [8]. We recently introduced an accurate and efficient multi-scale theory (atomic → monomer → filament) that overcomes some of these limitations when investigating electrical signal propagation along single wild-type actin filaments in physiological conditions [9]. The electrical signal that characterizes ion wave packets was shown to propagate as a monotonically decreasing soliton [10]. For the range of voltage stimuli and electrolyte solutions typically present in intracellular and in vitro conditions, the approach was able to predict a lower electrical conductivity with higher linear capacitance and non-linear accumulation of charge under the intracellular conditions. Additionally, the results showed a significant influence of the voltage input on the electrical impulse shape, attenuation, and packet propagation velocity. The filament sustained the soliton propagation at almost constant velocity for in vitro conditions, whereas intracellular conditions displayed a remarkable deceleration. Furthermore, the solitons were narrower and traveled faster at higher voltage input. As a unique feature, the multi-scale theory is able to account for the molecular structure conformation and biological environment changes often present in pathological conditions.

In this work, we present a Mathematica notebook, [11] which utilizes an extension of the multiscale approach, for an in depth study of monovalent ion wave packets traveling along single actin filaments in a variety molecular conformations and biological environments (https://github.com/MarceloMarucho/SignalPropagationPathologicalCondition). The program performs on single computers at very low computational cost, and does not require specialized training in computational methods, which can often be an obstacle for many students, researchers, and even experts in the field. Specific analyses can be achieved by making changes to the physiological and pathological conditions when selecting the desired nucleotide state (ATP or ADP), isoform (alpha-smooth, alpha-skeletal, alpha-cardiac, gamma-smooth, gamma-cytoplasmic, beta-cytoplasmic), and conformation model (wild-type or mutant) of the actin monomer. As a distinctive characteristic, a series of interactive plots are generated to elucidate the impact of pH, voltage input, and temperature on the filament conductivity, electric potential, velocity, peak attenuation and ionic wave packet profile. Overall, the program is fully editable. This feature allows users to change the default electrolyte aqueous solution and filament model parameters provided in the notebook.

In the following section, we describe the changes introduced on the multi-scale approach to study pathological conditions. In section 3, we describe the electric charge model of an actin monomer for different molecular conformations. Section 4 includes the organization, design and implementation of the interactive program. In section 5, we present an illustrative example, while section 6 includes the summary. A general overview of the multi-scale approach applied to a single (alpha-smooth muscle) wild-type actin filament is presented in the appendix. The explicit details of the theory can be found in a preceding article [9].

## 2. A multi-scale model for pathological conditions and the interactive program

The formulation of the multi-scale approach includes: (1) the atomistic scale; accounts for the pH of the aqueous solution, deprotonation of active residues on the protein, and the ionic composition of the biological environment, (2) the monomeric scale; models the ionic layering next to the charged protein, as well as, resistance to flow of ions in both the radial direction and along the filament, and (3) the filament scale; determines the velocity, acceleration, and soliton profile, along with the maximum travel time and distance of an ionic wave using a transmission line prototype model.

### 2.1. Biological solution model

#### 2.1.1. Aqueous solution

The pathological and physiological intracellular temperature gradient has been shown to range from 308*K* to 315*K* in the cytoplasm [12], around 1º*C* higher in the nucleus (when compared to the cytoplasm) [13], and around 321*K* at the mitochondria [14]. The interactive notebook allows users to choose a temperature between 298*K* and 320*K*. In this work, we account for the temperature dependence of the viscosity *μ* by using the temperature dependent equation *log*(*μ*) = −4.5318 − 247/(140 − *T*), where *T* represents the temperature in Kelvin and *μ* the viscosity in *Kg/m.s* [15].

The relative permittivities for the intracellular and in vitro conditions are set to 80*ϵ*_0_ and 78.358*ϵ*_0_, respectively, were *ϵ*_0_ is the permittivity of free space.

#### 2.1.2. Electrolyte

In this work, the dynamic mobility *u* for each ion species is estimated using the temperature dependent function *u*(*T*) = *u*_0_exp[−*U_o_/K_b_T*] [16, 17]. We obtained the parameters *u*_0_ and *U_o_* using the *FindFit* Mathematica function [18] and the experimental mobility values for several temperatures [19, 20]. This and other ionic species properties are shown in table 1. For instance, in figure 1a we show the fitting function for the sodium mobility for intermediate temperatures values between 275*K* and 320*K* (solid blue curve) and the corresponding experimental values (red dots).

**Table 1:**
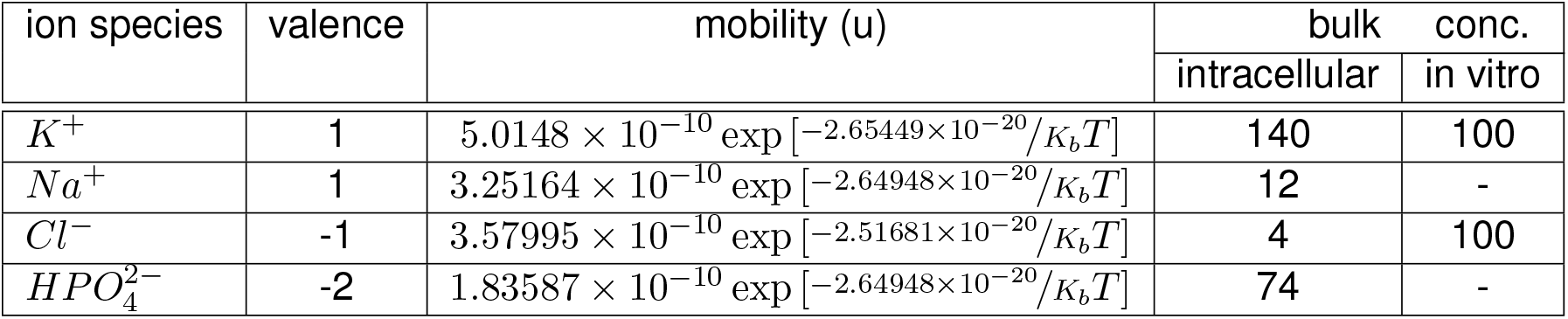
Ionic species used in both the intracellular and in vitro conditions. The last two columns represent the default concentrations used in the multi-scale approach for intracellular and in vitro conditions. The mobility is in units of *mol.m*^2^/*J.s*, and *K_b_* and *T* represent the Boltzmann constant and temperature in Kelvin. The unit for the bulk concentration is *mol/m*^3^.

| ion species | valence | mobility (u) | bulk | conc. |
| --- | --- | --- | --- | --- |
|  |  |  | intracellular | in vitro |
| $K^+$ | 1 | $5.0148 \times 10^{-10} \exp[-2.65449 \times 10^{-20}/K_b T]$ | 140 | 100 |
| $Na^+$ | 1 | $3.25164 \times 10^{-10} \exp[-2.64948 \times 10^{-20}/K_b T]$ | 12 | - |
| $Cl^-$ | -1 | $3.57995 \times 10^{-10} \exp[-2.51681 \times 10^{-20}/K_b T]$ | 4 | 100 |
| $HPO_4^{2-}$ | -2 | $1.83587 \times 10^{-10} \exp[-2.64948 \times 10^{-20}/K_b T]$ | 74 | - |

**Figure 1:**
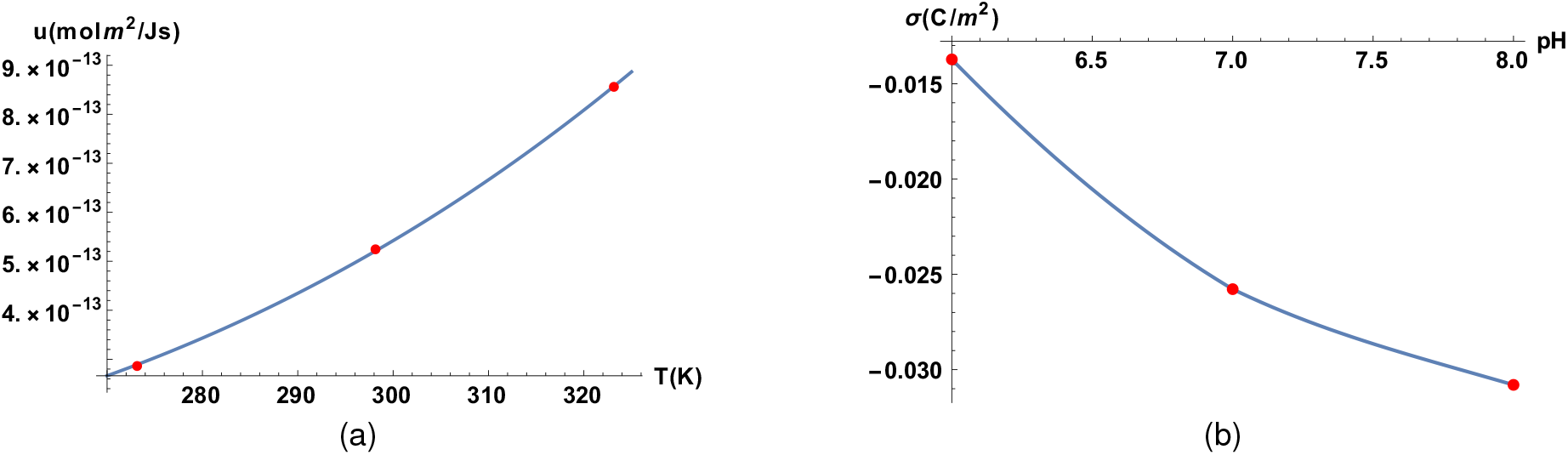
Validation on the Mathematica Interpolation functions. (a) The solid line represents sodium mobility for intermediate temperatures values between 275*K* and 320*K*. While, the red dots stand for the corresponding values obtained from experiments. (b) The solid line represents the surface charge density interpolating function for intermediate pH values between pH 6.0 and pH 8.0, whereas, the red dots are the values tabulated in table 1.

**Figure 2:**
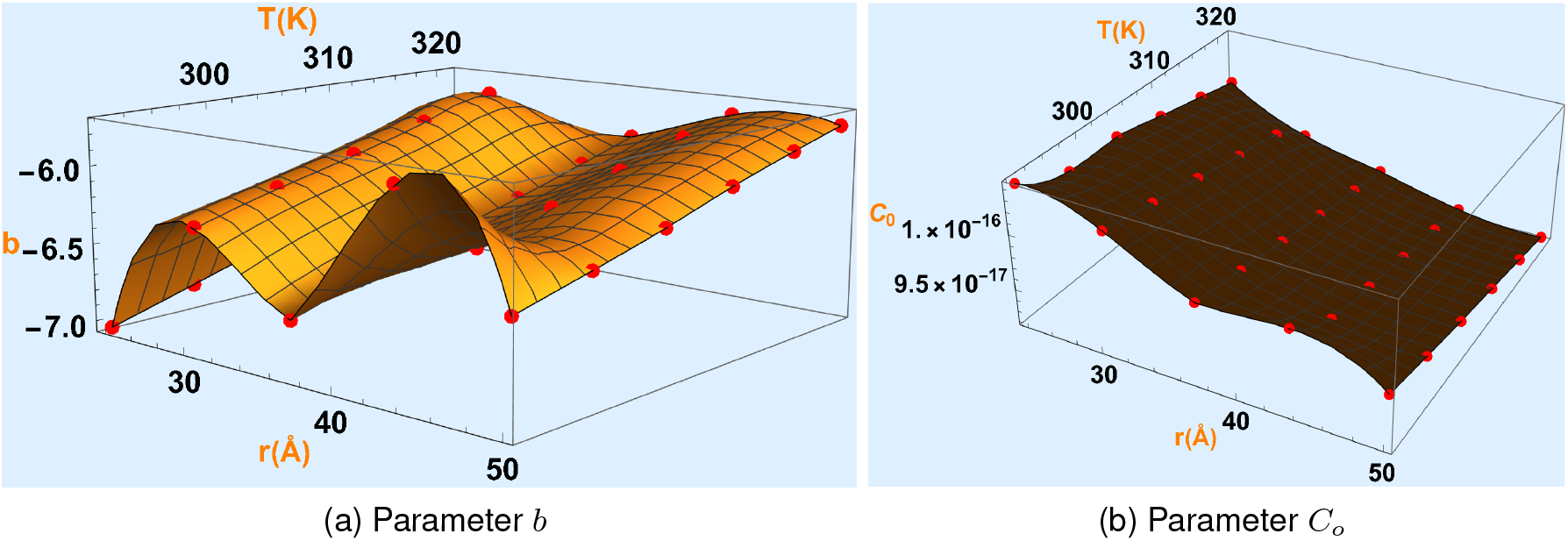
Capacitance interpolating surface functions for intracellular condition.

**Figure 3:**
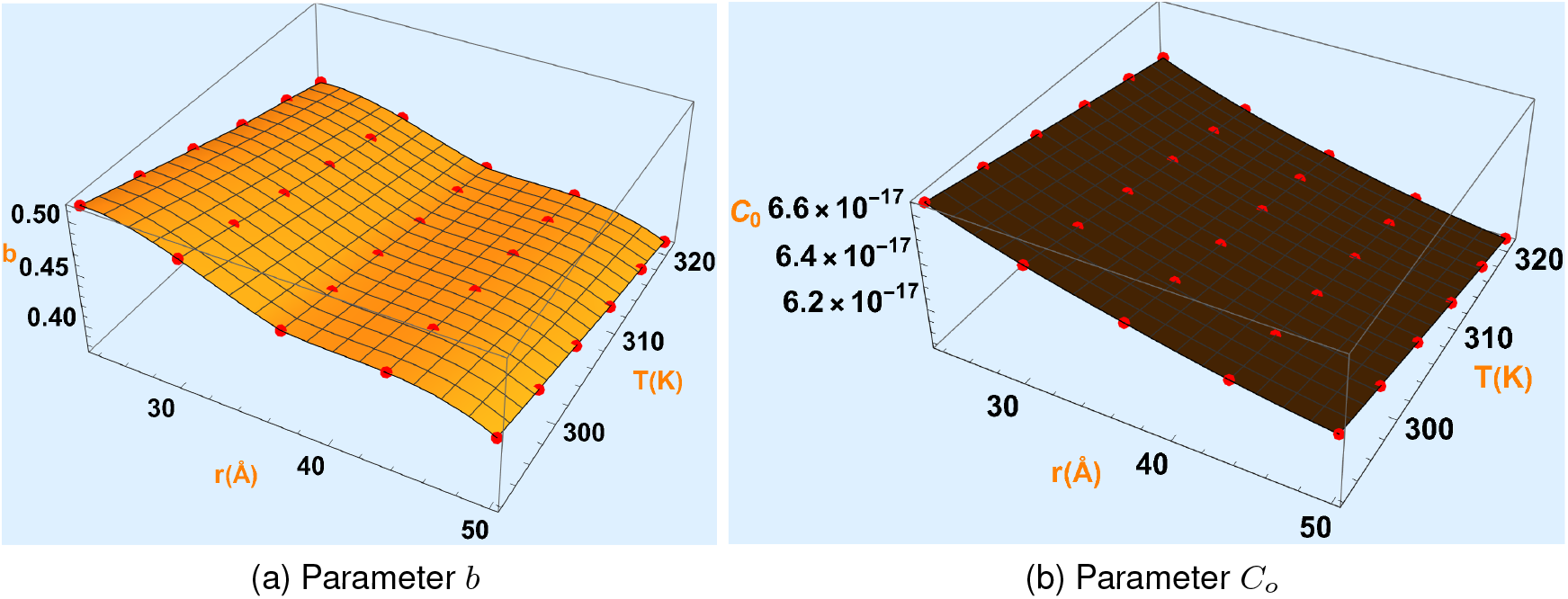
Capacitance interpolating surface functions for in-vitro condition.

### 2.2. Filament model

In table 2 (details are provided in the appendix) we tabulated the electric charge and surface charge density of the Cong molecular structure model [21] for the alpha wild type actin filament using the ATP state in physiological conditions. The intracellular pH for healthy cells lies between 6.8 and 7.3. On the other hand unhealthy cells, such as those infected with cancer, have been seen with a pH value between 7.2 and 7.8. Additionally, Alzheimer’s disease (AD) cells have shown a decrease in pH value when compared to healthy cells [22], being 6.5 the lowest pH found in brain ischemia [23]. As a unique feature, the surface charge density for intermediate pH values in the range between 6.0 and 8.0 were accurately estimated using the Mathematica Interpolation function [18], which uses third order fitting polynomial curves between successive data points. For illustration purposes, we display the discrete surface charge density values coming from the molecular structure and those coming from the interpolating function in figure 1b.

**Table 2:** The filament surface charge density (SCD) and electric charge obtained from the Cong (https://www.rcsb.org/structure/3B5U) molecular structure model for pH values 5-9, using the ATP state for a single (alpha-smooth muscle) wild-type actin filament. The electric charge was calculated using pdb2pqr webserver with an Amber force field (https://server.poissonboltzmann.org/pdb2pqr).

| pH | 14 monomers<br>filament charge<br>[$e$] | surface charge<br>density $\sigma$ [$C/m^2$] |
| --- | --- | --- |
| 5 | 23 | 0.006 |
| 6 | -82 | -0.021 |
| 7 | -154 | -0.039 |
| 8 | -184 | -0.047 |
| 9 | -222 | -0.056 |

### 2.3. Monomer model

The actin monomer length (*l*) is set to 5.4*nm*. The Mathematica interpolation function was used with surface charge density values to account for the pH dependence of the electric potential, electrical conductivity, resistances and impedance.

A Java Application for Cytoskeleton Filament Characterization (JACFC) web application [24, 25, 26] was used to obtain the capacitance parameters for different temperatures and filament radii.

A Mathematica interpolation was used to generate interpolating functions for the nonlinear parameter *b* and capacitance *C*_0_ for temperatures ranging from 292.15K to 321.15K, and filament radii ranging from 23.83Å to 50Å. Surface interpolating functions with JACFC data points are displayed in the plots 2 and 3 for intracellular and in vitro conditions, respectively.

## 3. Electric charge models for the nucleotide states, isoforms, and missense mutations

In the following subsections, we outline the method for determining the electric charge model for the nucleotide states, isoforms, and missense mutations. We also state the options available in the interactive program when choosing a mutation. A summary of the models is provided in table 3. More information on disease associated with actin, as well as a large list of phenotypes, genes and amino acid mutations can be found in the article by Parker et al. [27].

**Table 3:**
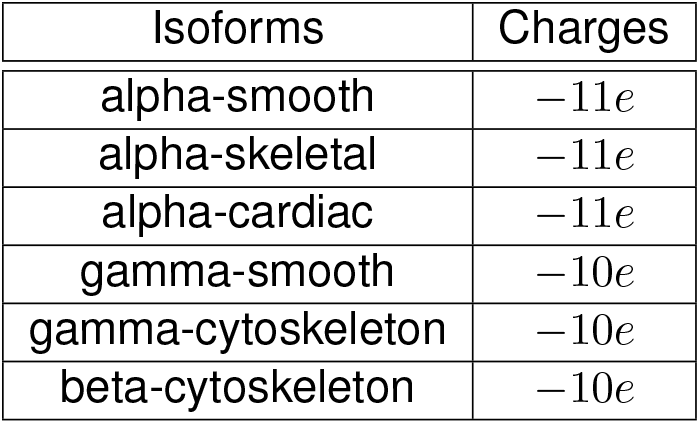
Electric charge models for actin protein isoforms, where *e* = 1.6 × 10^−19^*C* is the fundamental charge unit.

**Table 4:** Examples of charge changing mutations.

| isoform (gene) | mutation | charge | disease |
| --- | --- | --- | --- |
| skeletal α-actin (ACTA1) | Glu4Lys | 2 | Nemaline myopathy |
|  | Arg254His | -1 | DCM |
| smooth α-actin (ACTA2) | Gly36Arg | 1 | Thoracic aortic aneurysms and dissections |
|  | Arg38His | -1 | Thoracic aortic disease, coronary artery disease |
|  |  |  | coronary artery disease |
|  |  |  | strokes |
| β-actin (ACTB) | Glu362Lys | 2 | Neutrophil dysfunction and recurrent infection |
|  | Arg194His | -1 | Baraitser-Winter syndrome |
| cardiac α-actin (ACTC1) | Glu99Lys | 2 | HCM |
|  | Arg312His | -1 | DCM |
| γ-actin (ACTG1) | Glu314Lys | 2 | Hearing loss |
|  | Arg252Trp | -1 | Baraitser-Winter syndrome |
| smooth γ-actin (ACTG2) | Arg146Ser | -1 | Visceral myopathy, familial |

### 3.1. Nucleotide states and isoforms

From our pdb to pqr caculations for Cong wild type alpha-skeletal actin model we found the net negative charge per monomer is −11 electron charge. Based on previous studies, we assume that the surface charge density of the alpha wild type filament falls approximately 9% (1/11) when a *P_i_* (phosphorus ion) is released from the wild type actin filament following hydrolysis of the actin-bound ATP to ADP-*P_i_*. This results in a minus one charge difference between the ATP and ADP actin monomers. An in depth understanding of ATP and ADP actin states can be found in the work by Kudryashov et al. [28]. Similarly, we model gamma-smooth muscle, as well as gamma (*γ*) and beta (*β*) non-muscle (cytoplasmic) actin isoforms to have one less net negative charge than alpha (*α*) actin isoforms according to their amino acid sequence [29].

### 3.2. Missense mutations

The most common mutation in G-actin comes from missense mutations, which result in a substitution of a single amino acid on the wild-type actin monomer. The substitution on the mutated monomer may come in one of the following amino acid exchanges, where the symbol (q) is used to represent a charged amino acid residue that is affected by the pH of the solution (i.e. may lose (gain) a charge with an increase (decrease) in the pH level): (1) amino acid (q) residue → different amino acid (q) residue, (2) amino acid (q) residue → amino acid residue, (3) amino acid residue → amino acid (q) residue, or (4) amino acid residue → different amino acid residue. In cases (1) - (3), the type of residue removed and/or added, along with its charge at that pH value, are considered in the calculations of the new monomer surface charge density. On the other hand, in case (4) there is no change to the charge, because the amino acid side chain does not respond to the hydrogen concentration of the aqueous solution (i.e. pH of solution). These substitutions are represented using the three letter abbreviation of the amino acid being removed, the residue number, and the abbreviated name for the replacement amino acid (i.e. Abc#Def).

The ACTA1 mutations are recognized in diseases related to skeletal muscle, and have been estimated as 92% missense mutations [30]. Some consequences of these mutations were detected in hypertrophic cardiomyopathy (HCM), congenital myothopy, and muscular dystrophy. In this work we considered all the reported missense mutations in **(author?)** [27]. The user will be able to select a mutation from the drop down lists. The program will then compute the change in charge per monomer based on charged residues at the physiological pH range (pH 6 to pH 8). It is to note that mutations has effects on post translational modification, interaction with proteins, and change in charge. However, our approach can investigate any effect related to charge changing mutations. If a specific mutation does not effect charge we cannot expect any change in soliton propagation in this application. Some examples of charge changing mutations are shown in 4.

## 4. Results

### 4.1. Environmental changes due to Temperature

Using a more detailed description of the solvent viscosity for different temperatures suggest an investigation on how the temperature affects the effective electric conductivity, ion velocity profile, ion current density, and both the longitudinal and transversal ionic conductivities. For the invitro condition we compared room temperature (298.15*K*) with body temperature (310*K*). On the other hand, for the intracellular condition we compared body temperature (310*K*) with a high fever temperature (313*K*). One analysis (not presented here) showed negligible affects on the electric potential and ion concentration profiles due to the temperature changes. This was expected before hand, because there is only an indirect influence of the temperature which arises through Debye length. Additionally, the differences in temperature values used are rather narrow for biological conditions, that is a 3% increase for invitro and an even smaller 0.009% increase for the intracellular case. However, the effective electric conductivity between the actin surface and the bulk layer, as well as the ion current density distribution have noticeable changes. This is shown in figures 4 for the effective electric conductivity, and figure 5 for the ion current density results.

**Figure 4:**
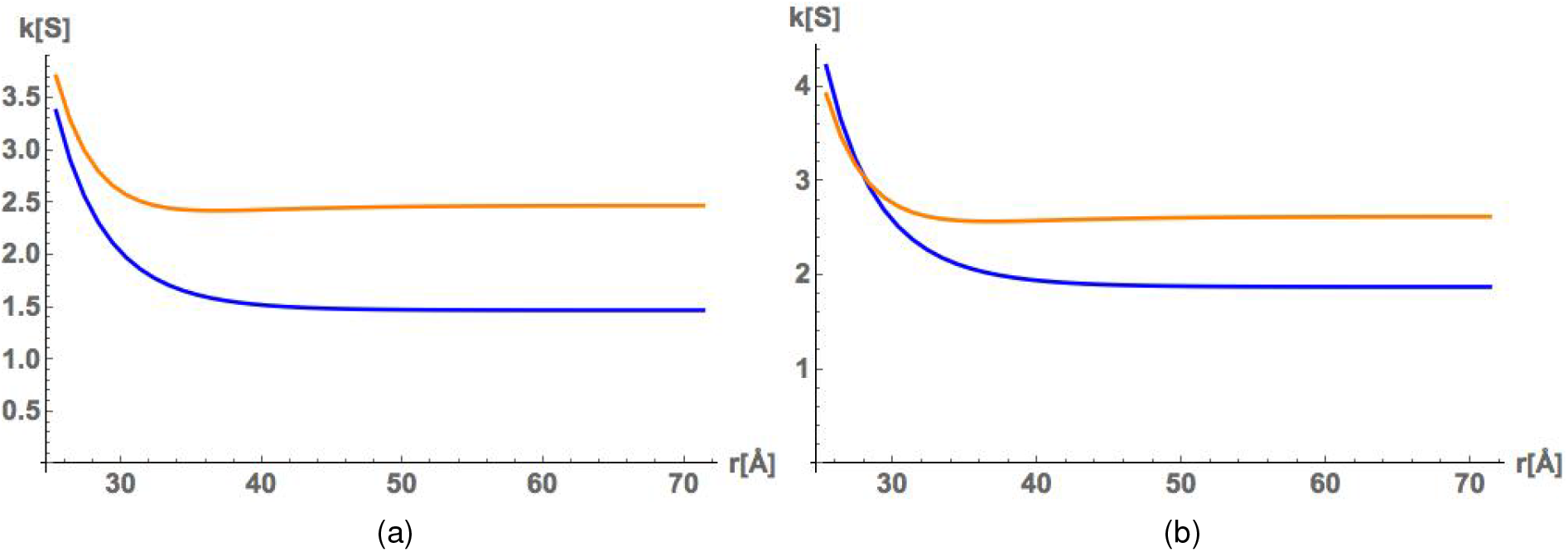
Temperature affects on the effective electric conductivity. The blue curves are for invitro conditions and the orange curves are for intracellular conditions. The radius is set to *R* = 23.83Å and the temperature is *T* = 298.15 (invitro) and *T* = 310*K* (intracellular) for subfigure (a) and *T* = 310*K* (invitro) and *T* = 313*K* (intracellular) for subfigure (b).

**Figure 5:**
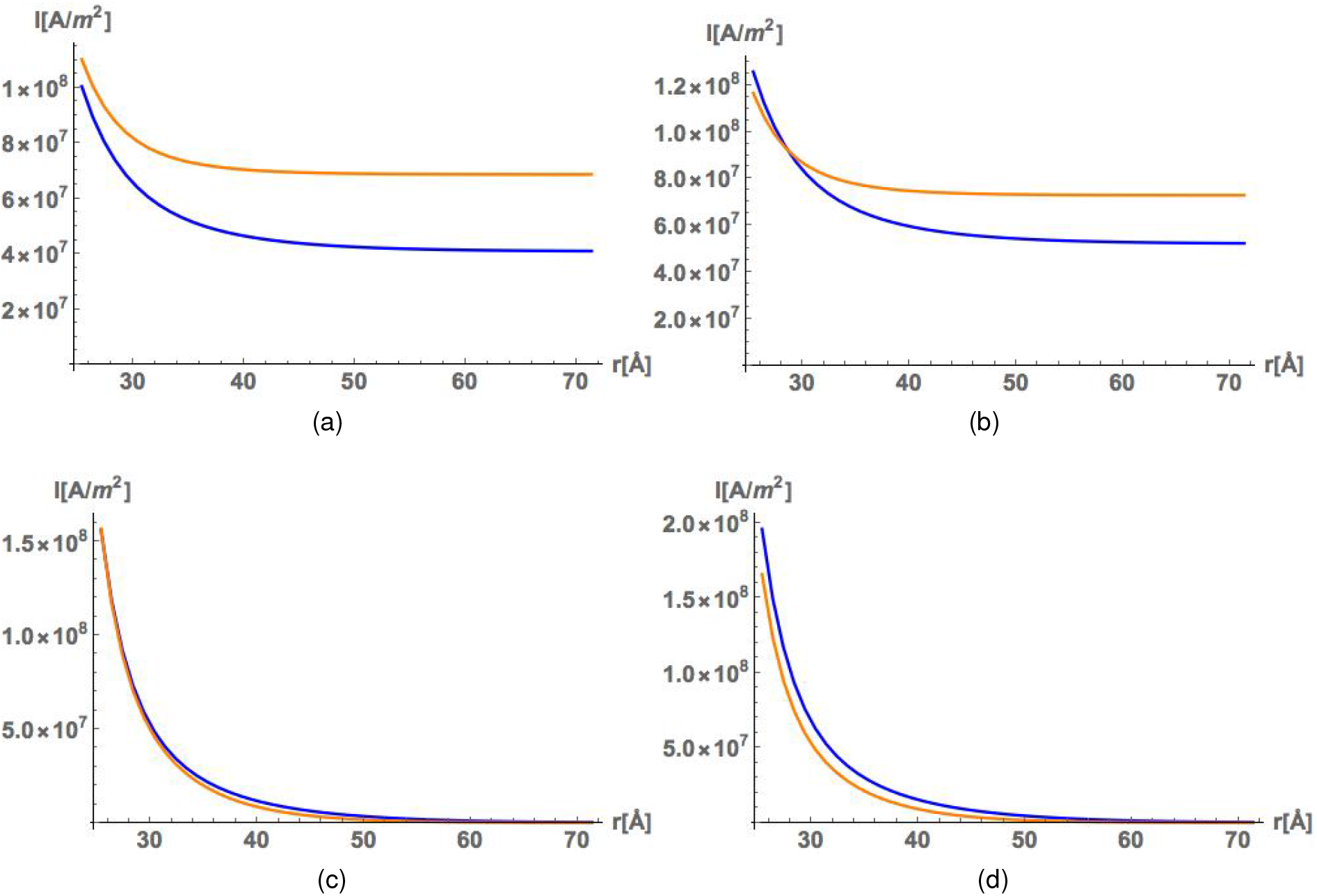
Temperature affects on the ion current density distributions in the radial direction starting at the actin surface. The blue curves are for invitro conditions and the orange curves are for intracellular conditions. Subfigures (a) and (b) represent the axial and radial directions, respectively, for temperatures *T* = 298.15 (invitro) and *T* = 310*K* (intracellular). Subfigures (c) and (d) represent the axial and radial directions, respectively, for temperatures *T* = 310*K* (invitro) and *T* = 313*K* (intracellular).

From the change in the conductivity there is a change in resistance due to their inverse relationship. The resistance has been demonstrated to play a role in the deceleration of the time averaged soliton velocity [Hunley et al.]. To better understand the transport of ionic information along actin filaments in different temperatures we looked at the velocity and peak attenuation of the traveling ionic soliton. The velocity is shown in figure 6 and the peak decay in figure 7.

**Figure 6:**
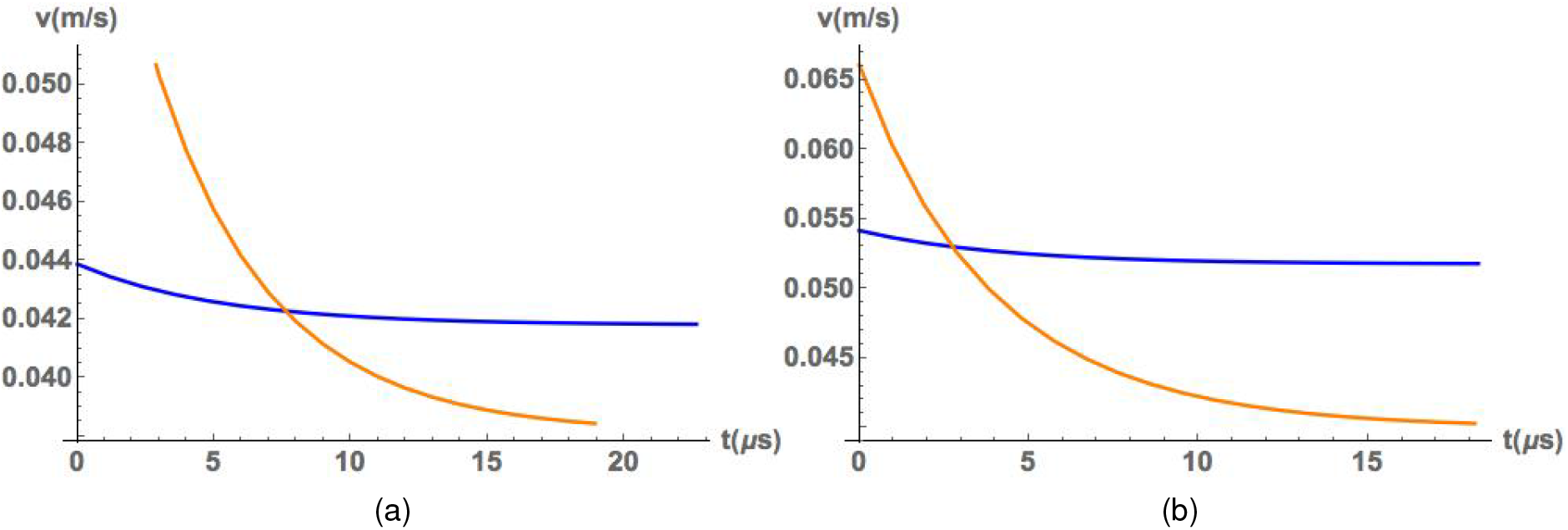
Temperature affects on the soliton velocity. The radius is set to *R* = 23.83Å with the blue curves used for invitro conditions and the orange curves for intracellular conditions. The voltage input is *V*_0_ = 0.15V, with the temperature at *T* = 298.15 (invitro) and *T* = 310*K* (intracellular) for subfigure (a) and *T* = 310*K* (invitro) and *T* = 313*K* (intracellular) for subfigure (b).

**Figure 7:**
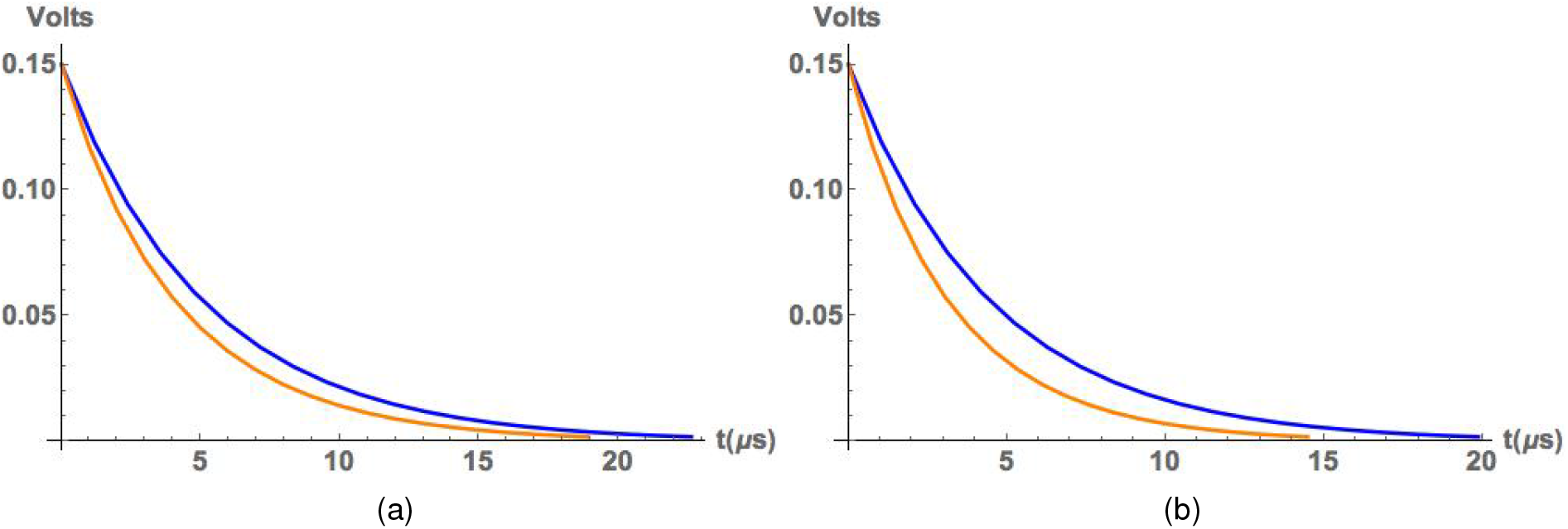
Temperature affects on the effective electric conductivity. The radius is set to *R* = 23.83Å with the blue curves used for invitro conditions and the orange curves for intracellular conditions. The temperature is *T* = 298.15 (invitro) and *T* = 310*K* (intracellular) for subfigure (a) and *T* = 310*K* (invitro) and *T* = 313*K* (intracellular) for subfigure (b).

### 4.2. Structural changes to the monomer

With a more accurate description of the diffuse layer coming from the non-linear behavior of the Poisson-Boltzmann distribution function, we look at the impact of the actin radius on the radial ion distribution. Using two different values for the radius, *r* = 23.83Å, 40Å, we compared the electric potential in figure 8.

**Figure 8:**
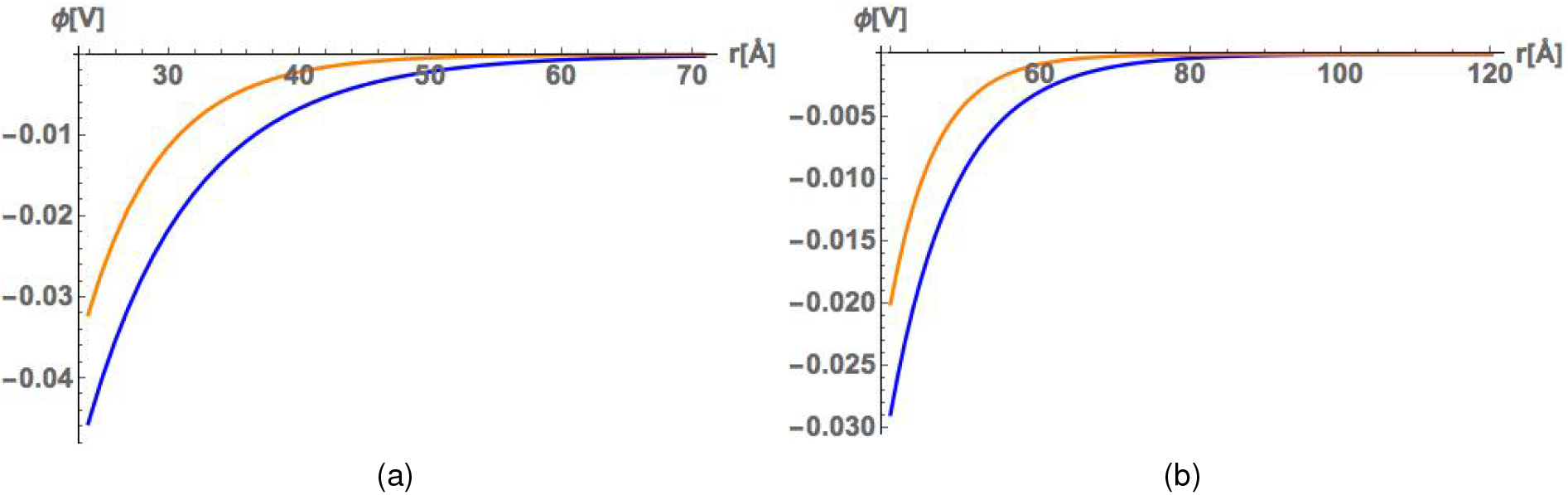
Impact of the actin radius on the electric potential profile. The subfigure (a) shows the electric potential with an actin radius of *R* = 23.83Å, whereas subfigure (b) uses a radius of *R* = 40Å. The blue curves are for invitro conditions with *T* = 298.15*K* and the orange curves are for intracellular conditions with *T* = 310*K*.

Different behaviors for the ion concentration in the radial direction starting at the actin surface are shown in figure 9 for different actin radius values, as well as changes to the effective electric conductivity (figure 10) and ion velocity (figure 11).

**Figure 9:**
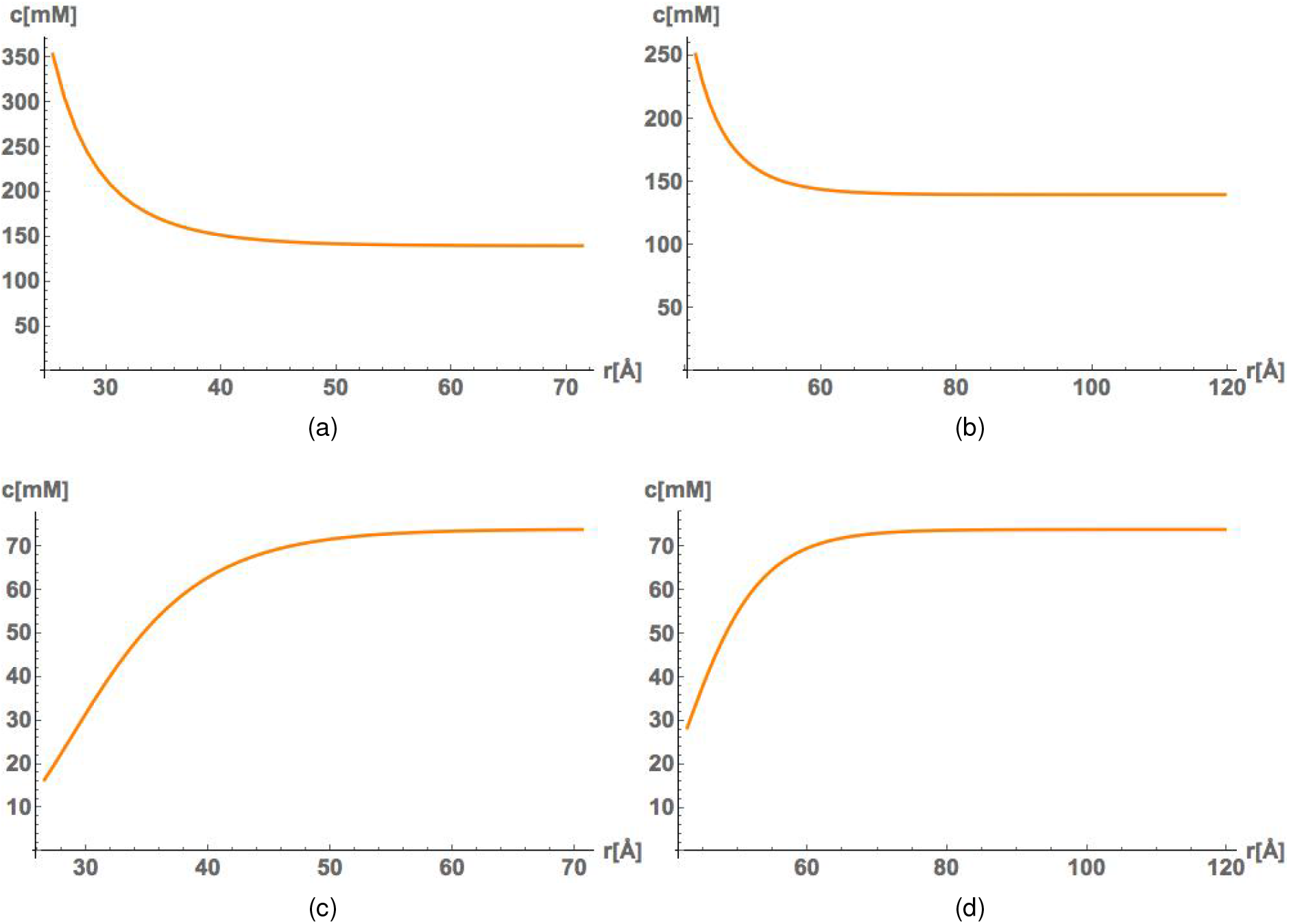
Impact of the actin radius on the ion distribution in the radial direction are shown for K^+^ in subfigures (a) and (b) for radius comparisons of *R* = 23.83Å and *R* = 40Å, respectively. In subfigures (c) and (d) we show the coin 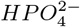 for *R* = 23.83Å and *R* = 40Å, respectively. The temperature is *T* = 298.15 for the blue curves showing invitro type conditions and *T* = 310*K* for the orange curves which represent an intracellular electrolyte mixture condition.

**Figure 10:**
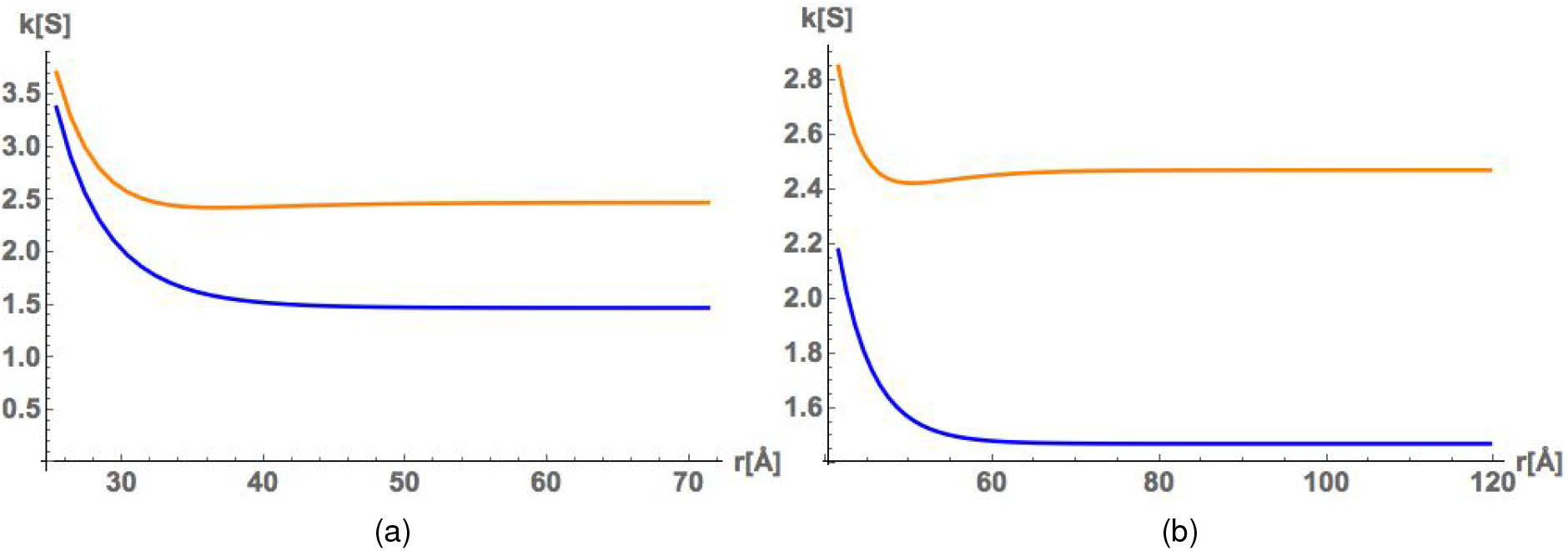
Impact of the actin radius on the electric conductivity profile. The blue curve is for invitro conditions with *T* = 298.15*K*, and the orange curve is for intracellular conditions with *T* = 310*K*. Subfigure (a) uses a radius of *R* = 23.83Å and subfigure (b) uses a radius of 40Å.

**Figure 11:**
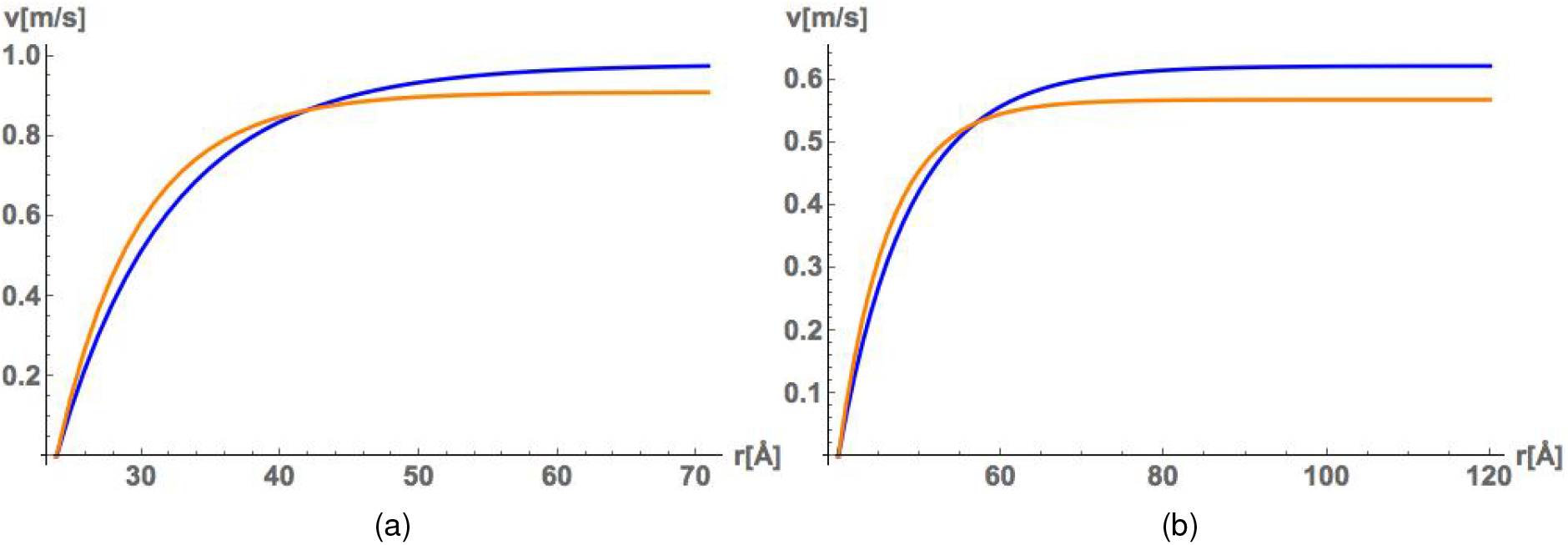
Ion velocity plots for invitro (blue) and intracellular (orange) electrolyte mixtures at temperatures *T* = 298.15*K* and *T* = 310*K*. Subfigure (a) is for 23.83Å and (b) is for 40Å.

The results of these changes on the soliton are represented in figure 12 by showing the velocity of the soliton wave packet for an actin radius of *R* = 23.83Å and *R* = 40Å. The temperature is *T* = 298.15 for the blue curves showing invitro type conditions and *T* = 310*K* for the orange curves which represent an intracellular electrolyte mixture condition.

**Figure 12:**
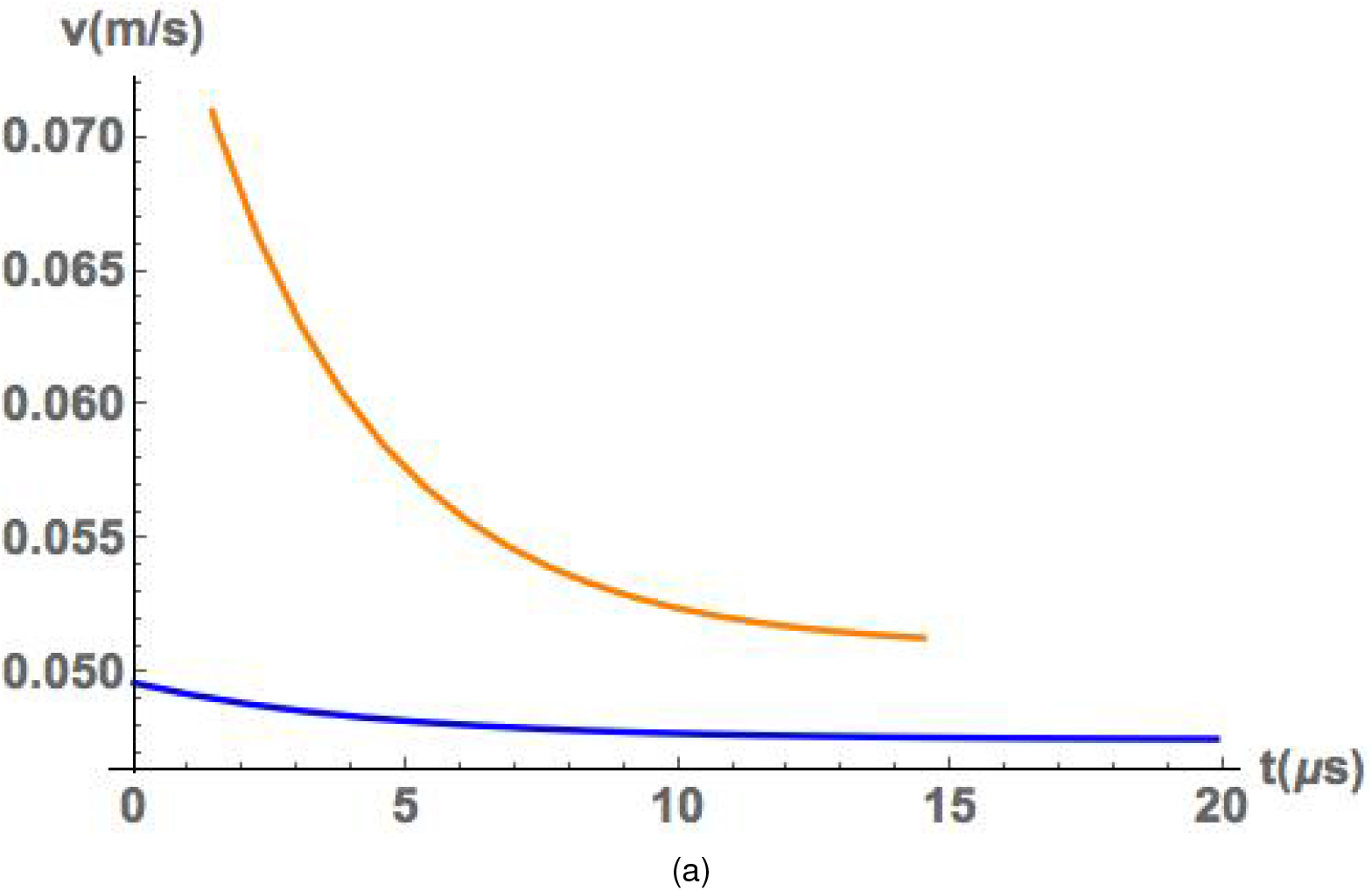
Impact of the actin radius on the velocity using a radius of *R* = 40Å. The temperature is *T* = 298.15*K* for the blue curve which represents invitro electrolyte concentration conditions, whereas the orange curve demonstrates intracellular electrolyte concentration conditions with *T* = 310*K*.

### 4.3. Environmental changes due to pH of the solution

Changes to the environment that result in different pH values of the surrounding solution have consequences to the ion concentration profiles for the intracellular condition. In figure 13 we show the results of the ion concentration profiles for the counterion *K^+^* and coion 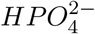 using a wild type actin monomer with a radius of *R* = 23.83Å for *pH* 6 and *pH* 8.

**Figure 13:**
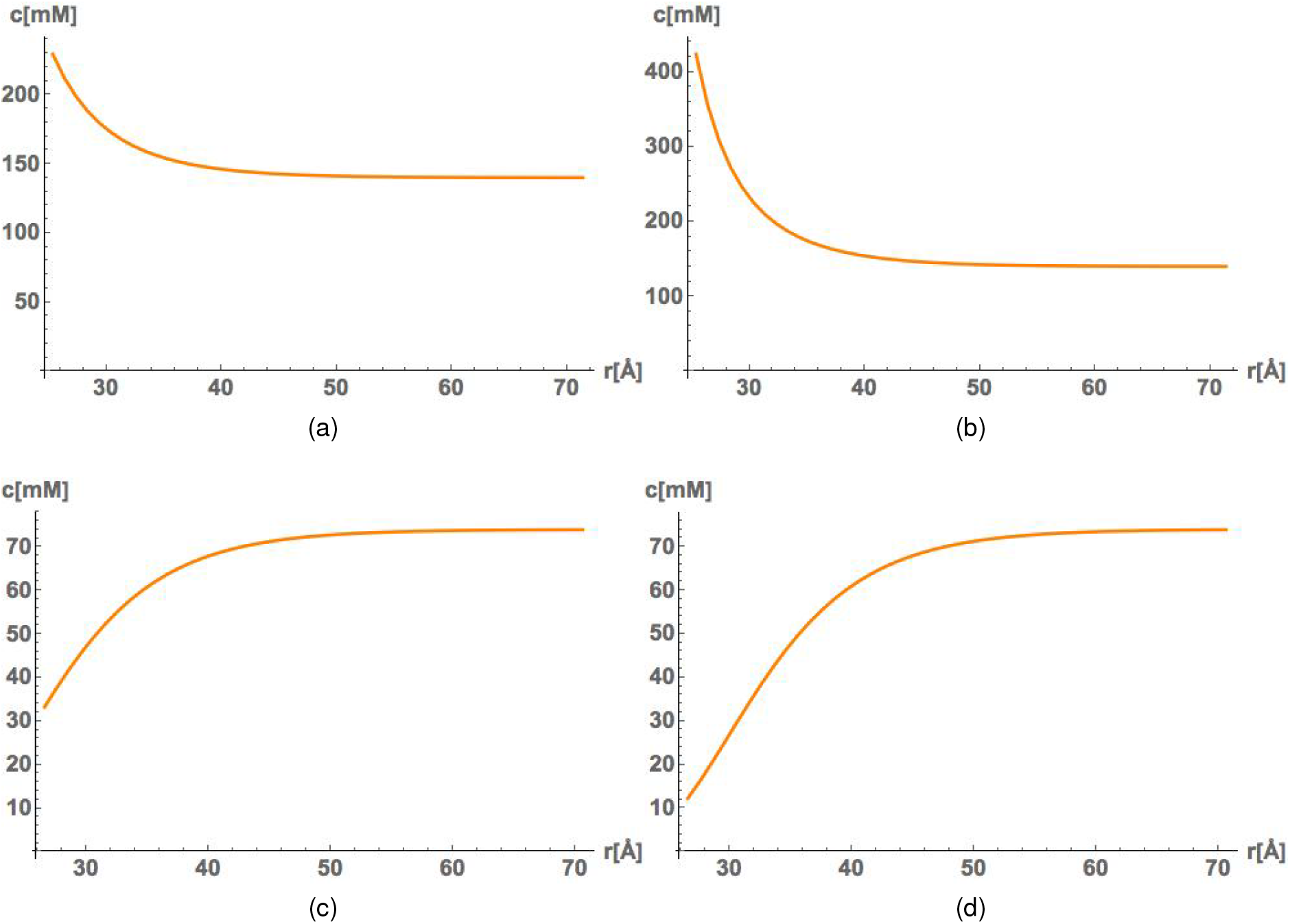
Ion concentration profiles for *K^+^* and 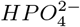 at two different pH values. In subfigures (a) and (b) we show the counterion distribution of *K^+^* for *pH*6 and *pH*8, respectively. In subfigures (c) and (d) we show the coion distribution of 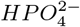 for *pH* 6 and *pH* 8, respectively. The independent variable on the horizontal axis starts at the radius of the actin monomer and extends in the radial direction toward the bulk layer.

The current density for *pH* 6 and *pH* 8 were also compared in figure 14, where the longitudinal (axial) profiles are shown in subfigures (a) and (b), and the transversal (radial) profiles are presented in subfigures (c) and (d).

**Figure 14:**
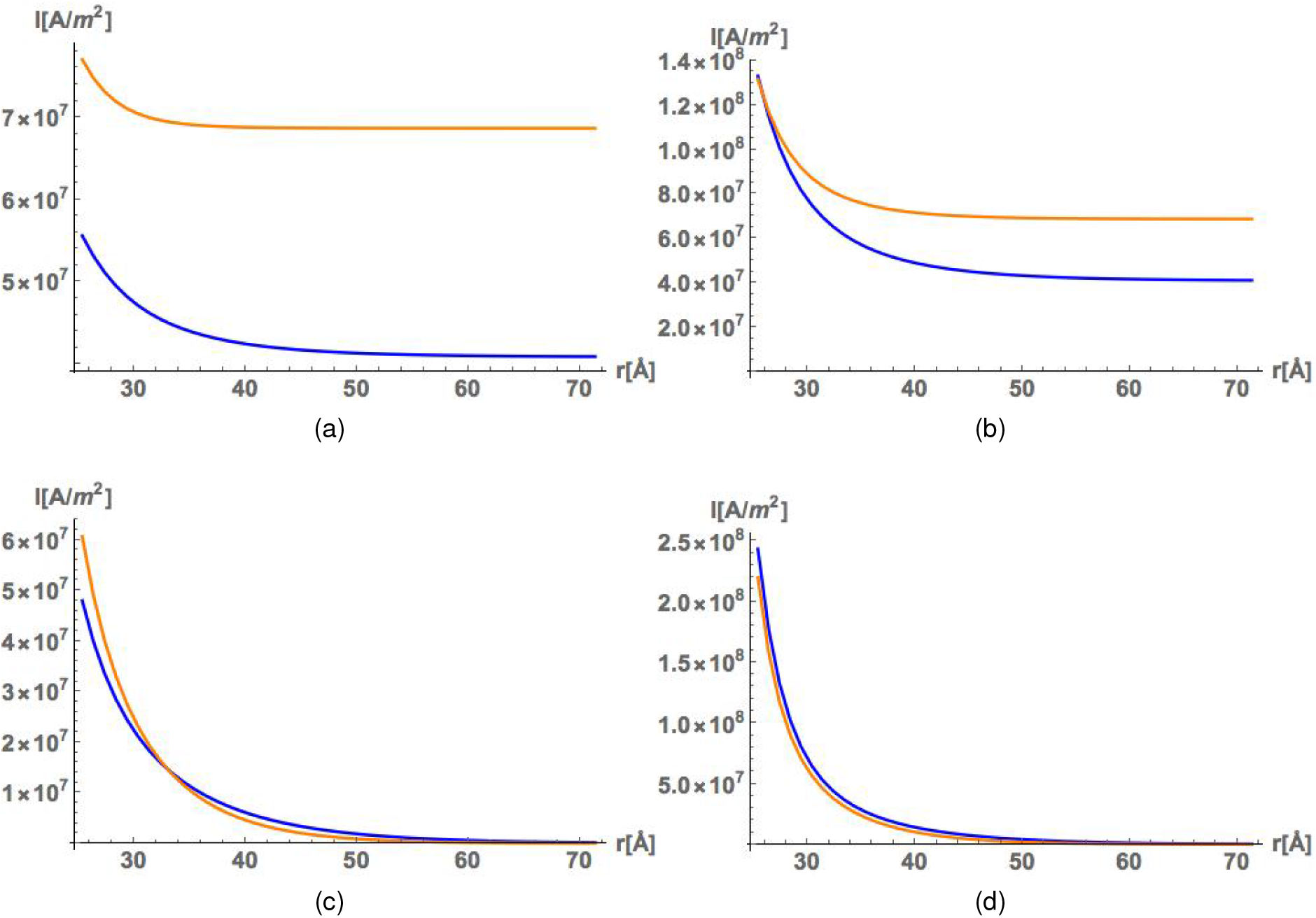
Current density profiles for the axial and radial directions at two different pH values using an actin radius of 23.83Å. The blue curves are for invitro conditions with temperature *T* = 298.15*K*, whereas the orange curves are for intracellular conditions with *T* = 310*K*. The subfigures (a) and (b) are for the axial direction with *pH* 6 and *pH* 7, respectively. The subfigures (c) and (d) are for the radial direction with *pH* 6 and *pH* 7, respectively.

### 4.4. Mutations

The high impact on the results coming from two different considerations for the radius in section 4.2 is due to changes in the surface charge density. In other words, an inversely proportional relationship results in a decrease in the surface charge density due to an increase in the actin radius. However, there is an alternative way to affect the surface charge density, which is by changing the charge of the actin monomer.

This is precisely the consequence of diseases that result in missense mutations. To analyze the impact of mutated actin monomers we choose an actin alpha-skeletal mutation (ACTA1), actin alpha-smooth muscle mutation (ACTB1), and an actin beta-cytoskeleton non-muscle mutation (ACTA2). Respectively, these correspond with nemaline myopathy (Glu4Lys); an actin monomer with 9e charges, falilial thoracic aortic aneurysm (Glu362Lys); an actin monomer with 8e charges, and Neutrophil dysfunction (Gly36Arg); an actin monomer with 12e charges. A comparison between ACTA2 and ACTB1 is seen using the electric potential in figure 15 and the ionic conductivity in figure 16.

**Figure 15:**
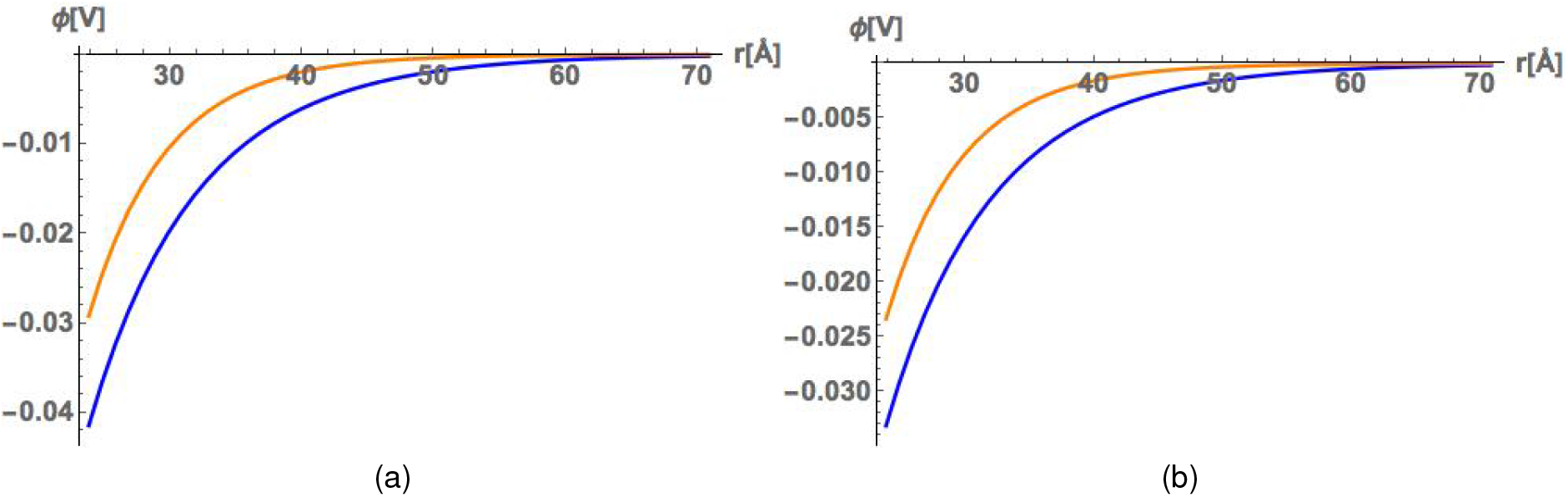
Impact of mutations on the electric potential profile. A comparison of the results on the electric potential response due to missense mutations from the isoforms *α*-smooth muscle (ACTA2) and non-muscle cytoplasmic (ACTB1) are shown. The mutation on ACTA2 is the amino acid residue replacement Gly36Arg in subfigure (a), whereas the ACTB1 is from the replacement of Glu362Lys in subfigure (b).The temperatures used in this analysis are *T* = 298.15*K* and *T* = 310*K* for invitro and intracellular electrolyte mixture conditions, respectively.

**Figure 16:**
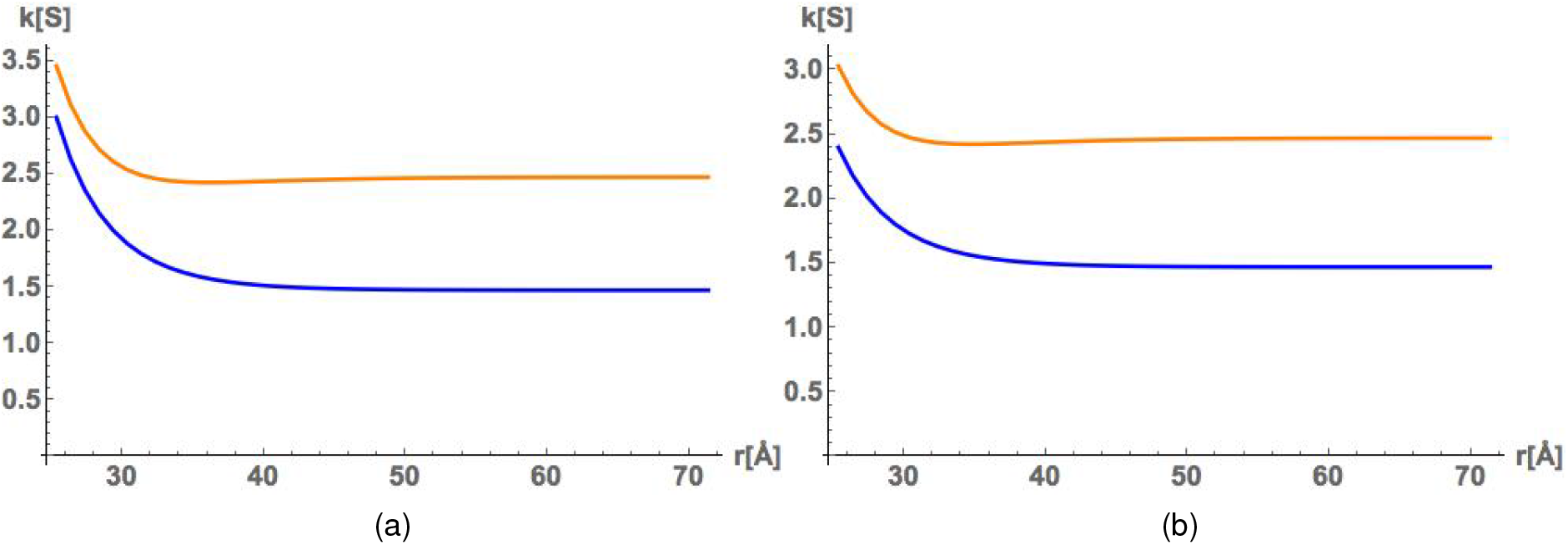
The affect of missense mutations on the ionic conductivity. The mutations from the isoforms *α*-smooth muscle (ACTA2) and non-muscle cytoplasmic (ACTB1) are shown. The mutation on ACTA2 is the amino acid residue replacement Gly36Arg in subfigure (a), whereas the ACTB1 is from the replacement of Glu362Lys in subfigure (b).The temperatures used in this analysis are *T* = 298.15*K* and *T* = 310*K* for invitro (blue) and intracellular (orange) electrolyte mixture conditions, respectively.

The diseased conditions resulted in different ion concentration profiles for the intracellular condition. This is demonstrated in figure 17, where we show the ACTA1 and ACTB1 results in plots (a) and (b), respectively, for K+ and HPO42-.

**Figure 17:**
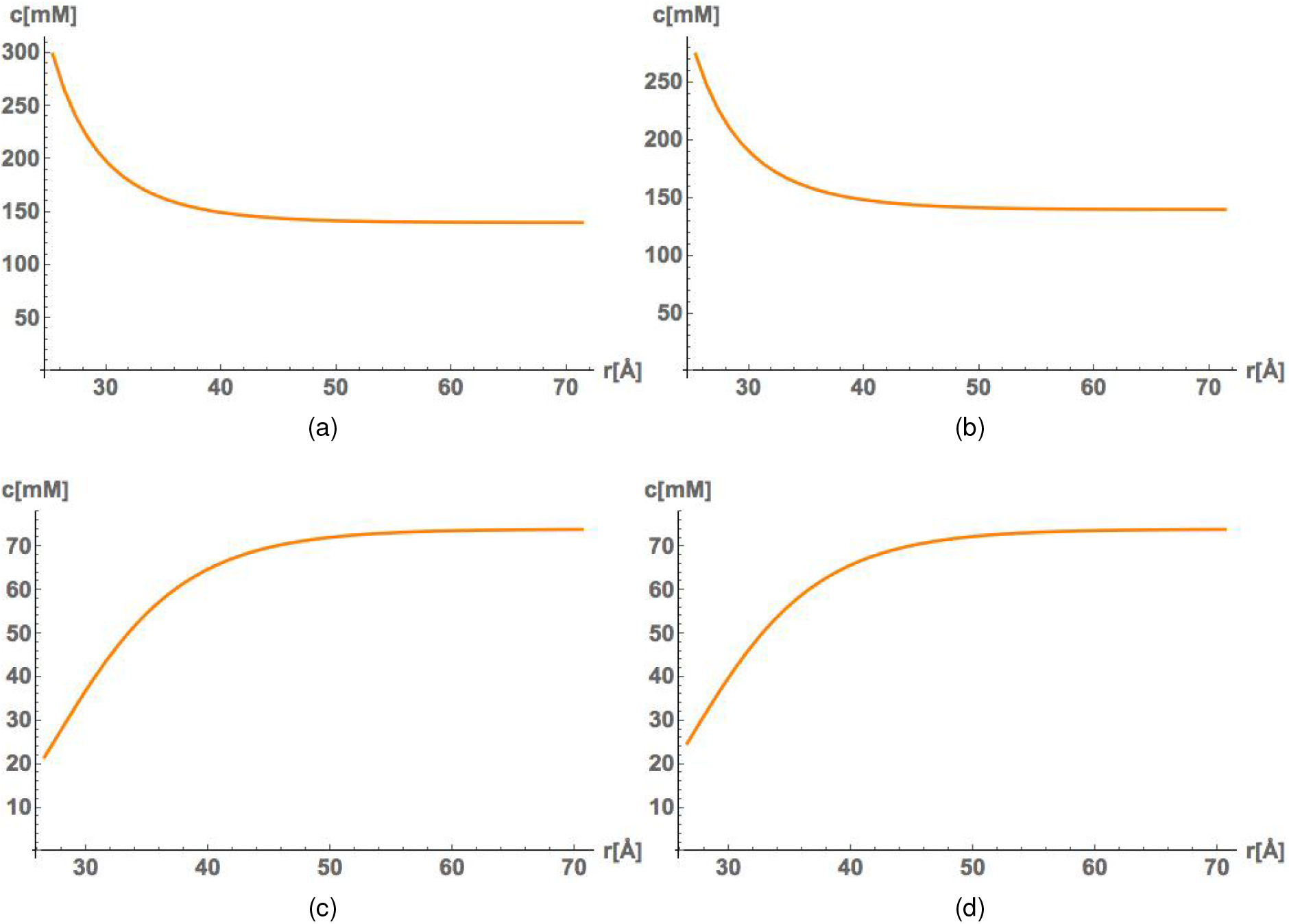
Mutation results for ion concentration distribution in the radial direction starting at the actin surface. The temperatures used in this analysis are *T* = 298.15*K* and *T* = 310*K* for invitro (blue) and intracellular (orange) electrolyte mixture conditions, respectively. The ions represented here are the counterion *K^+^* in plots (a) and (b) for a mutation to the isoforms ACTA1 and ACTB1, respectively, and the coion 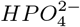 in plots (c) and (d) for a mutation to the isoforms ACTA1 and ACTB1, respectively.

The ion concentration profiles describe the way the ions are distributed in the diffuse layer. Results show counterionic species accumulate near the actin surface. Therefore, the counterions contribute to the larger conductivity along the filament, as confirmed by the ionic conductivity profiles in figure 16. This means there is a competitive relationship between the counter- and coions when it comes to conductivity contributions, which therefore, impact the resistance resulting in different soliton velocities (Fig. 18).

**Figure 18:**
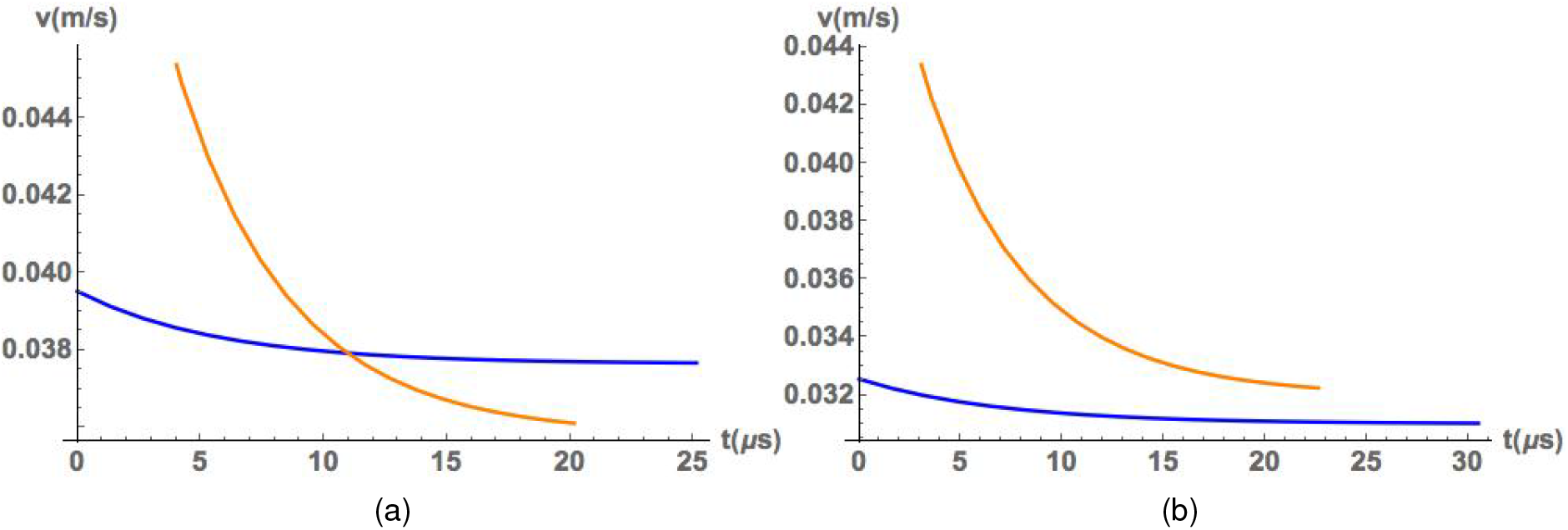
Impact of mutations on soliton velocity using the Gly36Arg and Glu362Lys missence mutations in subfigures (a) and (b), respectively. The temperatures used in this analysis are *T* = 298.15*K* and *T* = 310*K* for invitro (blue) and intracellular (orange) electrolyte mixture conditions.

## 5. Discussion

### 5.1. Temperature

The temperature changes for the values chosen in this work show a difference of Δ*T_invitro_* = 11.85*K* for invitro conditions and Δ*T_intracellular_* =3*K* for intracellular conditions. These values were used because an invitro experiment may measure at room temperature (298.15*K* = 25°*C*) and also at body temperature (310*K* = 36.85°*C*). The intracellular condition is a much smaller difference since we started at body temperature, and therefore, only increased to a high fever temperature of (313*K* = 39.85°*C*). Certainly, the larger change for invitro conditions has a more impressive impact in all results analyzed.

For the effective electric conductivity we see an increase due to increasing the temperature 4. Mathematically this comes from the Boltzmann distribution, where the temperature is in the denominator of the decreasing exponential term. Physically, the result is due to increased thermal motion causing a smaller restraining force on the couterions near the surface, and an intensified presence of the external field the ions. This is also visualized in figure 5 by the invitro results shown by the blue curves. The axial current density equation is analyzed by showing how the ion current density distribution contributes in different parts of the diffuse layer. Near the surface of the actin there is a very evident increase in the current density between figures 5(a) and 5(b) for the invitro results. For the intracellular results the smaller increase in temperature leads to a much smaller increase in the current density. The rise in the horizontal asymptote in both conditions shows the conductivity in the bulk layer is also affected by the temperature change.

The Bjerrum length measures the separation between counterions and coions due to the competition between the electrostatic interactions and the thermal energy. Since the Bjerrum length is inversely proportional to the temperature, a temperature increase results in a smaller separation (i.e. a more narrow electrical double layer). The results of this can be seen in the radial ion current distribution in figures 5(c) and 5(d). The increase in the radial current density throughout the diffuse layer suggest ions moving in toward the actin surface.

The traveling soliton is a packet of ions in the diffuse layer that travels along the actin filament. Therefore, the increase in conductivity corresponds to a decreased resistance, and therefore, a ionic soliton with a faster velocity. In figure 6 we show how the soliton velocity changes with increasing temperature. A comparison between the invitro and intracellular curves in figures 6(a) and 6(b) show a temperature increase does indeed result in a wave packet with a faster velocity. However, figure 7 suggest a faster steeper decay for a soliton with a larger velocity. Therefore, increasing the temperature will produce a faster moving soliton that may travel approximately the same distance as the soliton with a slower velocity.

### 5.2. Increase in actin radius

The electrostatic potential is proportional to the surface charge density of actin, which can be calculated from the linear charge density divided by the actin radius times 2*π*. That is, increasing the radius will decrease the surface charge density and also the magnitude of the electrostatic potential. This was the result when we compared a actin radius of *R* = 23.83Å in figure 8(a) with an actin radius of *R* = 40Å in figure 8(b). Additionally, we expect a smaller accumulation of counterionic species at the surface and a larger number of coionic species due to the reduced electrostatic forces. Figure 9 shows this assumption was correct. The larger radius contained a 28% decrease in *K^+^* ions at the surface, whereas the 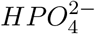 ions had a 86% increase.

The ionic concentration changes resulted in a drastic change to the electrical conductivity by resulting in a 21% decrease for the intracellular results (orange) curve and 35% decrease for the invitro results in figure 10. The smaller decrease in the intracellular conditions could be a result of the divalent coions 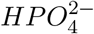 constraining some counterions in the region near the surface.

The results on the ion velocity have the expected consequences of a decreased potential, because the ion velocity is proportional to the potential difference of the location measured in the EDL and the slipping plane. This can be visualized in figure 11(a) and 11(b) by the decrease in the horizontal asymptote of the larger radius 11(b). On the other hand, the soliton velocity plots show an increase due to the larger radius (compare figures 6(a) and 12). This was counter intuitive because the ionic axial velocity was just argued to decrease. However, resistance is inversely proportional to cross-sectional area which means an increase in actin radius also decreases the resistance. Hence, there is a competitive relationship between the decrease in the axial velocity profile and the decrease in the resistance when it comes to the soliton velocity.

### 5.3. Changes to the pH of the solution

To investigate the impact of pH we considered the ion concentration profiles and current density profiles for the axial and radial directions. The consequence of pH changes on actin can be seen in table 2, where we see that a larger pH results in a more negative charge on the actin monomer in physiological conditions (pH 6-pH 8). Therefore, a stronger attraction was expected by counterions, along with a stronger repulsion for coions, with increasing pH values. This was confirmed in figures 13. A *K^+^* increase of 82% was found at the actin surface, whereas a 68% decrease was concluded for 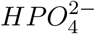. This can be correlated with a large increase in the longitudinal (axial) current density profile near the surface (14(a) and 14(b)). Unlike the results on current density for temperature changes there is no increase in conductivity for the bulk layer. Additionally, a larger electrostatic attraction implies a increase in couterion current density toward actin surface. This is seen in figures 14(c) and 14(d) for the radial current density profile comparison at pH 6 and pH 8.

### 5.4. Mutations

Missense mutation can result in change to the charge of an actin monomer. This comes from removal and replacement of an amino acid residue. Some mutations exchange amino acids that respond to the pH of a solution. For example, replacing glutamic acid (Glu) with a lysine (Lys) has a consequence to the monomer at physiological pH values. Therefore, we analyzed the electrostatic potential of the missense mutations Gly36Arg and Glu362Lys in figures 15. The Gly36Arg amino acid exchange is a ACTA2 isoform mutation that results in an actin monomer with a −12*e* charge 15(a). On the other hand, the Glu362Lys amino acid exchange is a ACTB1 isoform mutation that results in an actin monomer with a −8*e* charge 15(b). Hence, the ACTA2 isoform mutation in this case results in a stronger electrostatic response. Consequently, figure 16 shows a larger ionic conductivity coming from the ACTA2 mutation 16 (a) when compared to the ACTB1 mutation 16 (b).

We also compared a actin-skeletal muscle isoform mutation (ACTA1) with the actin non-muscle cytoskeletal isoform mutation (ACTB1). In this case, the amino acid exchange for the ACTA1 isoform was Glu36Lys, which results in a −9*e* actin monomer. A comparison of the ionic concentration distributions for *K^+^* and 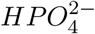 is represented in figure 17. These figures show the expected result of more couterions and fewer coions approaching the actin surface for the ACTA1 iso form mutation (i.e. for the Glu36Lys amino acid exchange).

The analysis on velocity profiles for the traveling ionic wave packet are seen in figure 18. A mutation resulting in a larger negative charge show a faster traveling wave 18(a), when compared to a actin monomer with a smaller charge 18(b). This is a result of the increased conductivity (figure 16) causing a lower resistance. Other analysis can be performed with different mutations in the interactive Mathematica notebook outlined in the appendix.

## 6. Conclusion

Due to their conducting bionanowire properties, actin filaments have been experimentally demonstrated and theoretically modeled to be capable of transmitting ions across their filament length when a Gaussian input voltage is applied to the filament. The solitons produced by the transmission line model represent ionic waves, which travel distances in the neighborhood of a micron, and could be a biological consequence in the local vicinity of cells [31].

In this work, we used a non-linear Boltzmann description of the ionic distribution in the electrical double layer to perform a detailed and accurate analysis on the potential profile, ion velocity, effective electric conductivity, ion concentration profile, ion current density, soliton velocity, and amplitude attenuation of a ionic soliton wave. We studied environmental changes such as temperature affects and pH changes of the surrounding solutions, as well as, structural changes to an actin monomer due to radius changes. Additionally, we investigated for the first time the electrostatic consequences of actin mutations from different disease conditions.

Temperature changes and pH differences, which are known to occur in unhealthy muscle and non-muscle cells, were shown to result in different ion accumulations at the surface of the actin monomer (and filament), ionic conductivities and ionic soliton wave packet velocities. The changes in radius showed a competition between the ion velocity profiles and the resistance when influencing the soliton propagation velocity.

This study found that a comparison of the pH changes and the mutation analysis both resulted in common trends. Specifically, there were impacts to the ion concentration profiles, current density, and traveling ionic solitons. This is because pH fluctuations in the solvent change the charge of the protein as explained above. That is, table 2 showed the charge of the Cong unit is −83e, −154e, and −184e, for pH values 6, 7, and 8, respectively. To be explicit, that is approximately −6e, −11e, and −13e charges per monomer for increasing pH. In the analysis on mutations, we used −8e, −9e, and −12e for the mutated actin monomeric isoforms ACTB1, ACTA1, and ACTA2 (see section 3 for how these calculations were made). This leaves an open question regarding the connection between the pH value in the intracellular space of cells and diseases involving missense mutations with at least one of the seven pH dependent amino acid. Despite a missense mutation only replacing one amino acid on an actin monomer, the results on the ionic wave packet velocity show a consequential outcome.

Additionally, we introduced a novel interactive Mathematica notebook that allows for the characterization of the soliton propagation along actin filaments for a wide range of physiological and pathological conditions. In particular, the user has the option of selecting a nucleotide state, isoform of interest and its associated disease. As a unique feature, the investigator may perform their own detailed study on the conductivity, mean electric potential, velocity, and peak decay by using an interactive slider for adjustments of the pH (6.0–8.0), and voltage input (0.05*V*–0.40*V*). This simple and easy to use program requires no specialized training or expertise in the field, and is designed to conduct a detailed analysis on both wild-type and mutated conformations. The simplicity of the theoretical formulation, and the high performance of the Mathematica software, enable the analysis of multiple conditions without computational restrictions. These studies may provide an unprecedented molecular understanding on how actin filaments are related to aging and disease, due to a more complete understanding of their electrical properties in both muscle and non-muscle cells [31]. A future version of this program will account for the propagation of divalent ionic waves along actin filaments.

## APPENDIX

### A. Design and implementation

The organization of the Mathematica notebook is presented in Figure 19. In section 1 on the notebook, we included and prcessed all the reported mutations in **(author?)** [27]. In section 2 and 3 we defined the electrolyte aqueous solution and filament model parameters described in sections 2 and 3 on this article. While section 4 of the program contains the soliton equations corresponding to the transmission line theory for actin filaments outlined in the appendix. Finally, the interactive plots are provided in section 5 on the notebook.

**Figure 19:**
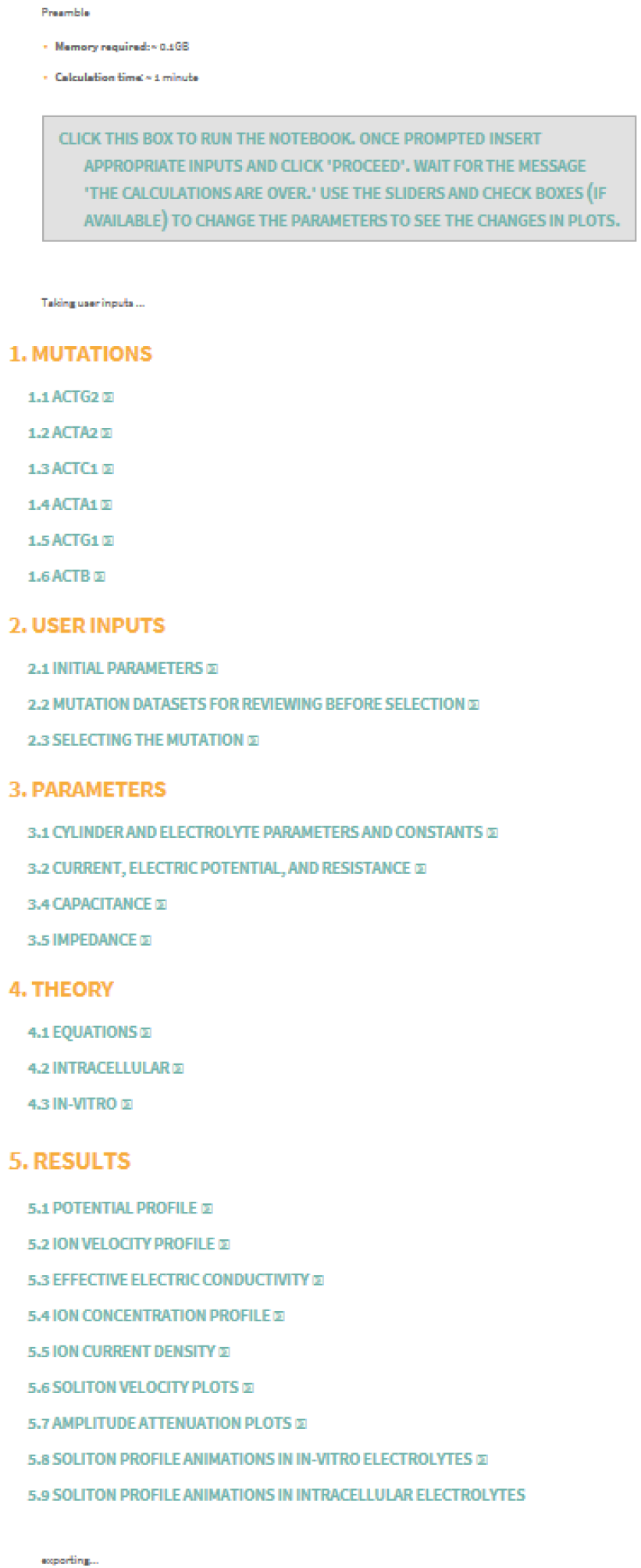
Mathematica notebook content organization

The initiation of the program is started by clicking the “run notebook” box located at the top of the interactive Mathematica notebook (Figure 20). The program allows for simple to use input parameters (radius and surrounding temperature), along with the choice of actin ATP or ADP assembled monomers, six choices of actin isoforms, and either a wild-type or mutated actin conformation (see Figure 21). If the mutant option is selected, two additional windows will open, one will show all the mutations along with related disease name to review and the other will provide a drop down list of all the muations to selecet from. (see Figure 22). After the program has completed all computations and rendering of all plots, the last screen will open to notify the user that the calculations are finished (see Figure 23). A text file named actin_input.txt will be downloaded to the current directory containing key input parameters for the run 24. For several runs the text files will be numbered.

**Figure 20:**
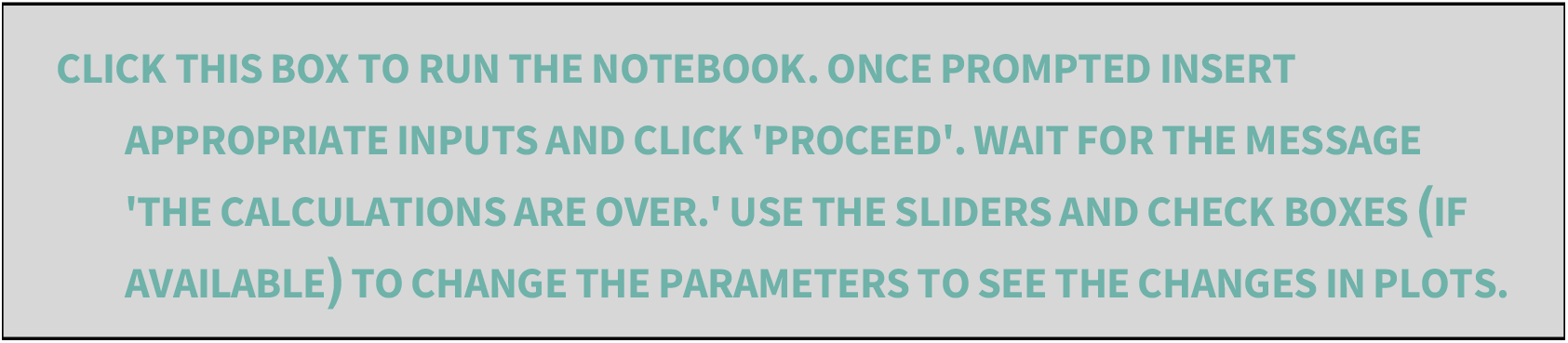
Starting the program: Upon opening the Mathematica notebook the user has to click in the box where it states “Click this box to run the notebook… .”

**Figure 21:**
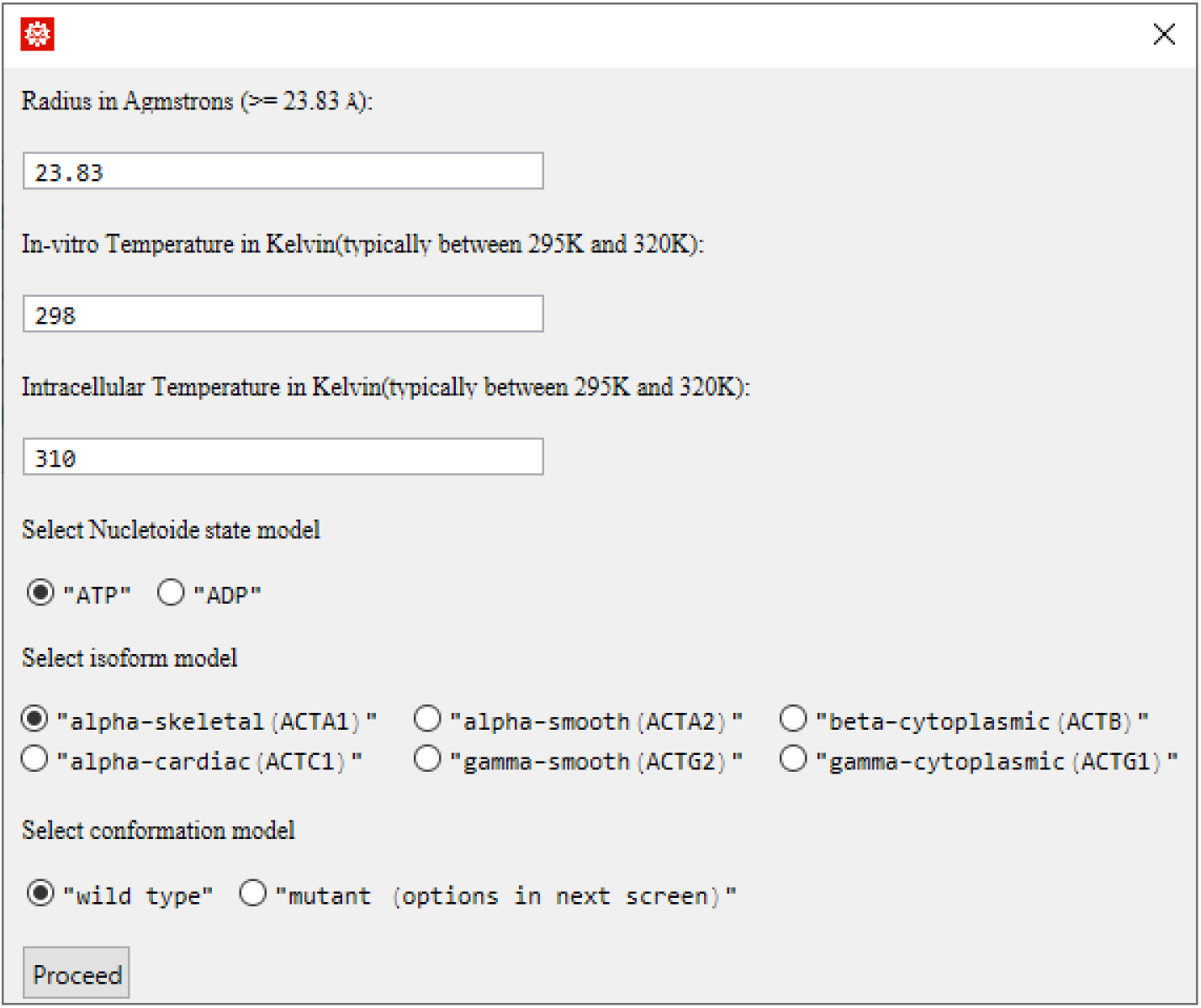
After clicking the “run notebook” box, a dialog box will open and ask the user to set the input parameters (Radius in Angstroms, Temperature in Kelvin, Nucleoide state model, isoform model, and conformation model)

**Figure 22:**
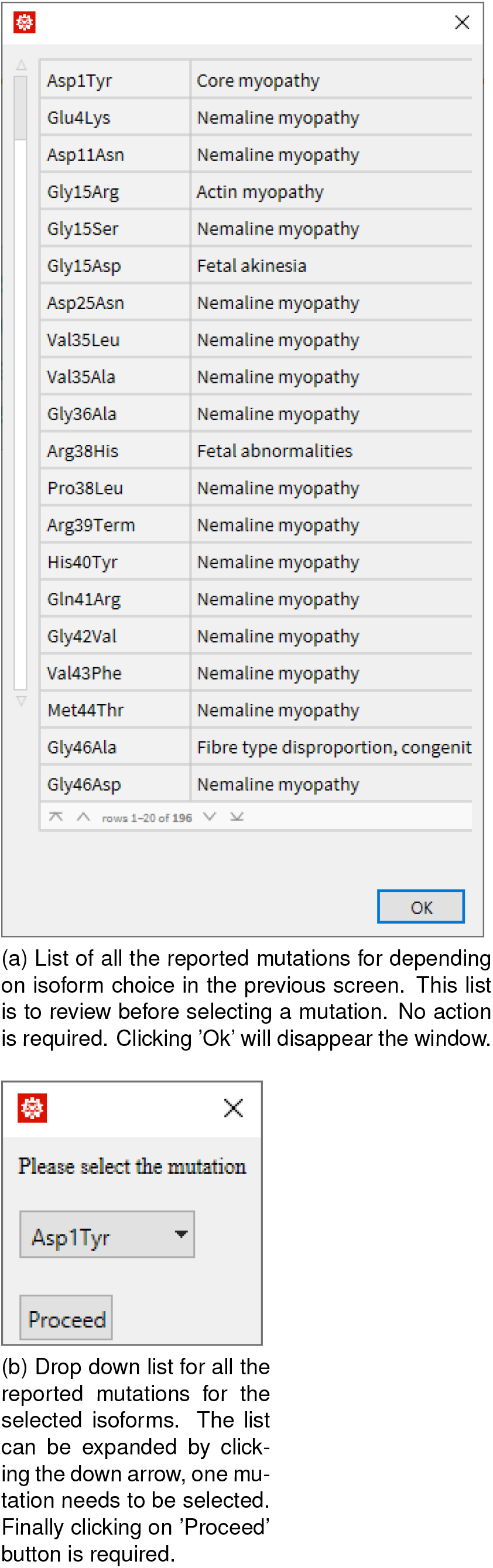
If the user selects the “mutant” option, two windows will open simultaneously, one to review all the reported mutations and the other to select a mutation from a drop down list.

**Figure 23:**
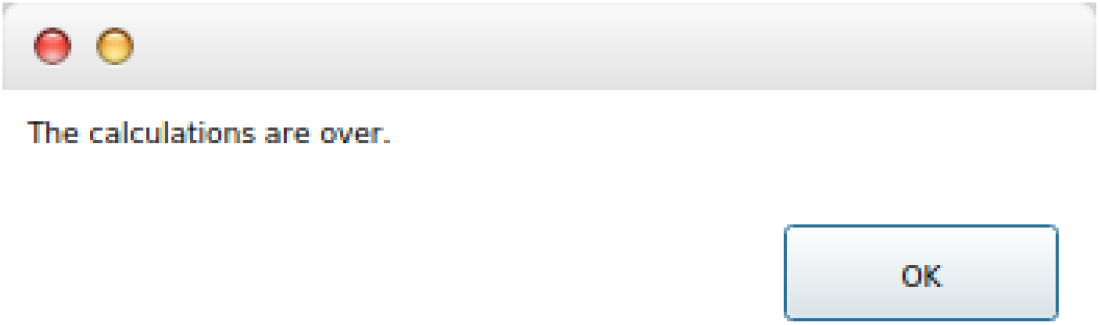
After the program completes all calculations and rendering all plots, a pop-up message will inform the user that “The calculations are over.”

**Figure 24:**
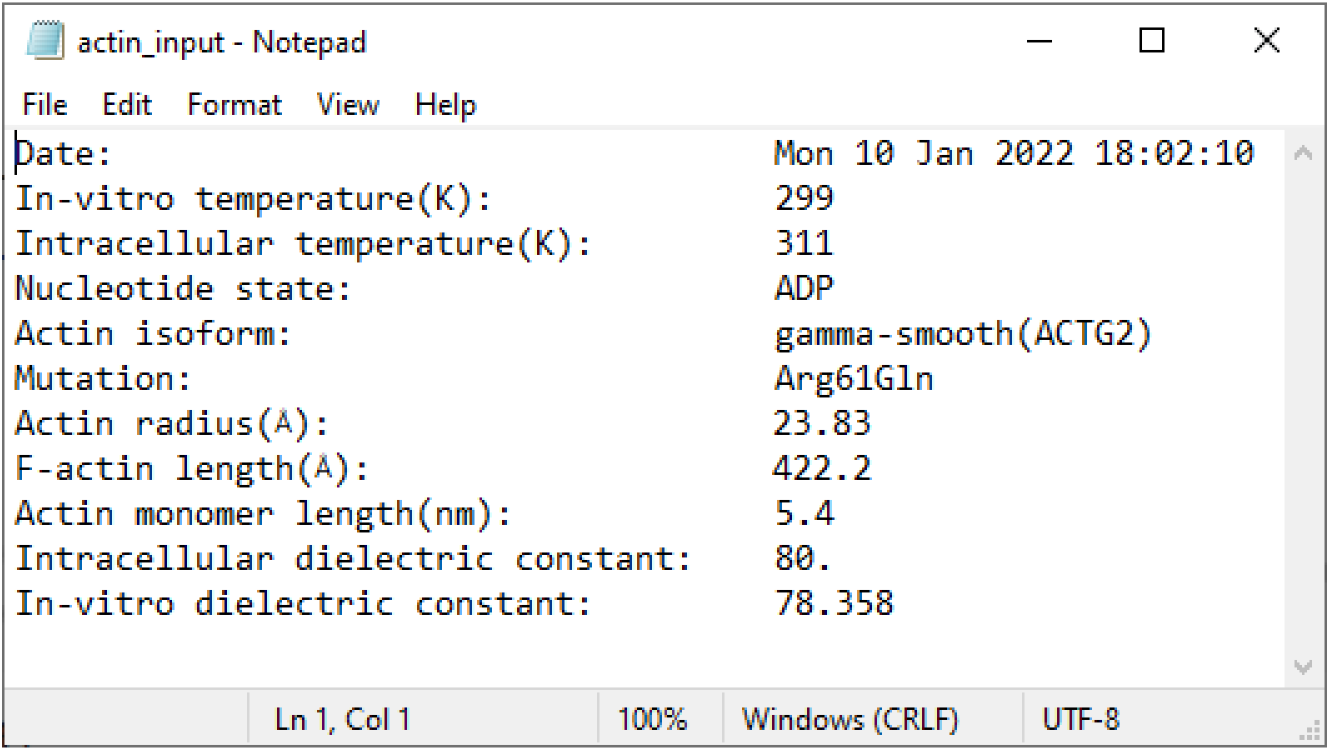
A text file will be saved in the current directory containing the key input parameters.

#### A.1 Interactive options

There are three important parameters that can be manipulated in the interactive plots to obtain a more detailed analysis. These are the following: (1) the condition of the biological environment (in vitro or intracellular), (2) the input voltage, and (3) the pH of the electrolyte solution. A brief explanation for each of the three options is given below.

##### A.1.1 In vitro and Intracellular conditions

All the results can be analyzed at both in vitro and intracellular conditions. In this program, the in vitro environment represents a surrounding aqueous electrolyte solution containing potassium chloride *KCl* as the electrolyte, and the intracellular model includes potassium chloride *KCl*, sodium (*Na*), as well as hydrogenphosephate 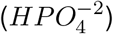 to represent larger charged anions in the biological environment (see table 1). For the velocity and amplitude attenuation plots, the in vitro and intracellular conditions can be selected independently, or chosen simultaneously. However, there are two sections for the soliton results that represent the in-vitro and intracellular conditions separately.

##### A.1.2 Voltage input

In some plots, the input voltage is being used to compare in vitro and intracellular conditions, while a pre-defined domain or range is used in others. In the first situation the input voltage cannot be adjusted by the user (see figures 28a and 31). Otherwise, the input voltage can be chosen using the slider for values between 0.05*V* − 0.4*V* volts. This voltage input range includes the values used in invitro experiments [32, 6]. For the soliton profiles, both in vitro and intracellular sections have an animation that compares two voltage inputs of 0.05*V* and 0.15*V* while the slider represents the change in time.

##### A.1.3 pH

The interactive notebook allows users to analyze pH solutions in the range between 6.0 and 8.0. A lower pH value comes from a greater number of hydroxide ions in the solution. This means more positively charged hydrogen ions protonate the residues on the actin monomer and decrease the charge of the filament when the G-actin is negative (see table 2). Otherwise, a higher pH decreases the number of hydroxide ions in solution due to deprotonating the active residues on the monomer, which results in an increase in the negative charge of actin. Thereby, pH solution affects the formation of the EDL due to fewer counterions accumulating at the filament surface.

##### A.1.4 Using the slider

The results in the Radial Electric Potential, Velocity, and Amplitude Attenuation sections include interactive plots with an adjustable slider that allows extensive analyses of the pH and input voltage. The sliders can be adjusted manually by sliding left or right, and also by clicking the “plus” sign at the end of the bar for finer adjustments (Figure 25). Additionally, a pH value can be fine tuned to a tenth of the value desired (Figure 25a, 25c), where as the voltage can be adjusted to two hundredths of the nearest whole number (Figure 25b, 25d).

**Figure 25:**
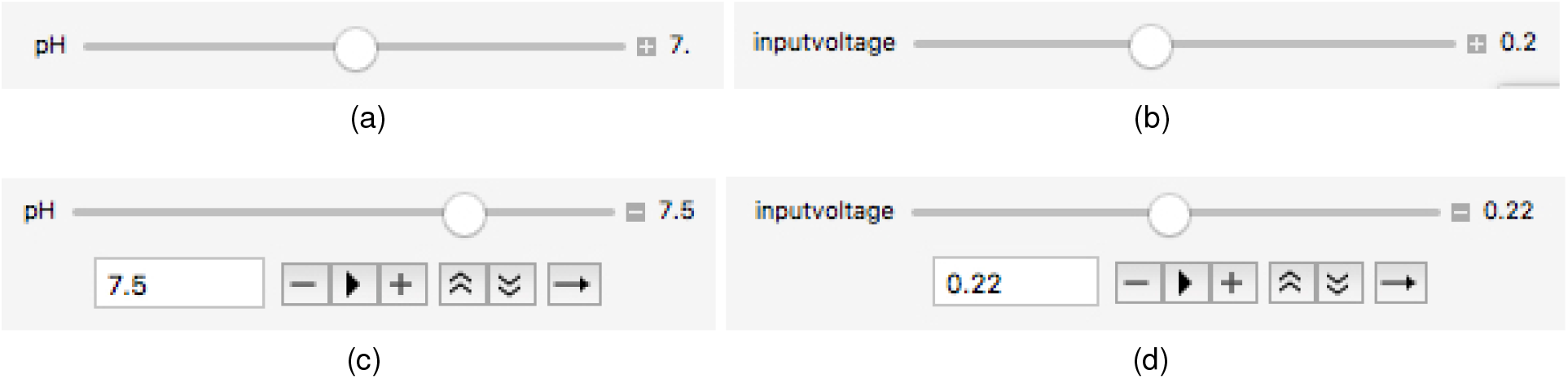
The slider for the respective plots can be changed to fine tune results for a specific analysis. In the sub plots, the slider has been adjusted between the discrete values of 7.0, 7.1, 7.2, 7.3, 7.4, and 7.5. for pH in plots (a) and (c), and 2.0 and 2.2 for in put voltage in plots (b) and (d).

### B. Illustrative example

For physiological conditions, the Mathematica program reproduces the results and plots provided in reference [9]. Below, we provide the results and a brief analysis for an actin filament of radius equal to 35Å in the ADP nucleotide state and immersed in a electrolyte solution at 310K.

#### B.1 Effective filament conductivity

Figure 26 shows the effective filament conductivity results as a function of the pH. It can be seen that the relative filament conductivity in a in vitro electrolyte aqueous solution (blue curves) is lower than in the intracellular condition ( orange curves). . This behavior was observed in invitro experiments with actin filaments [32].

**Figure 26:**
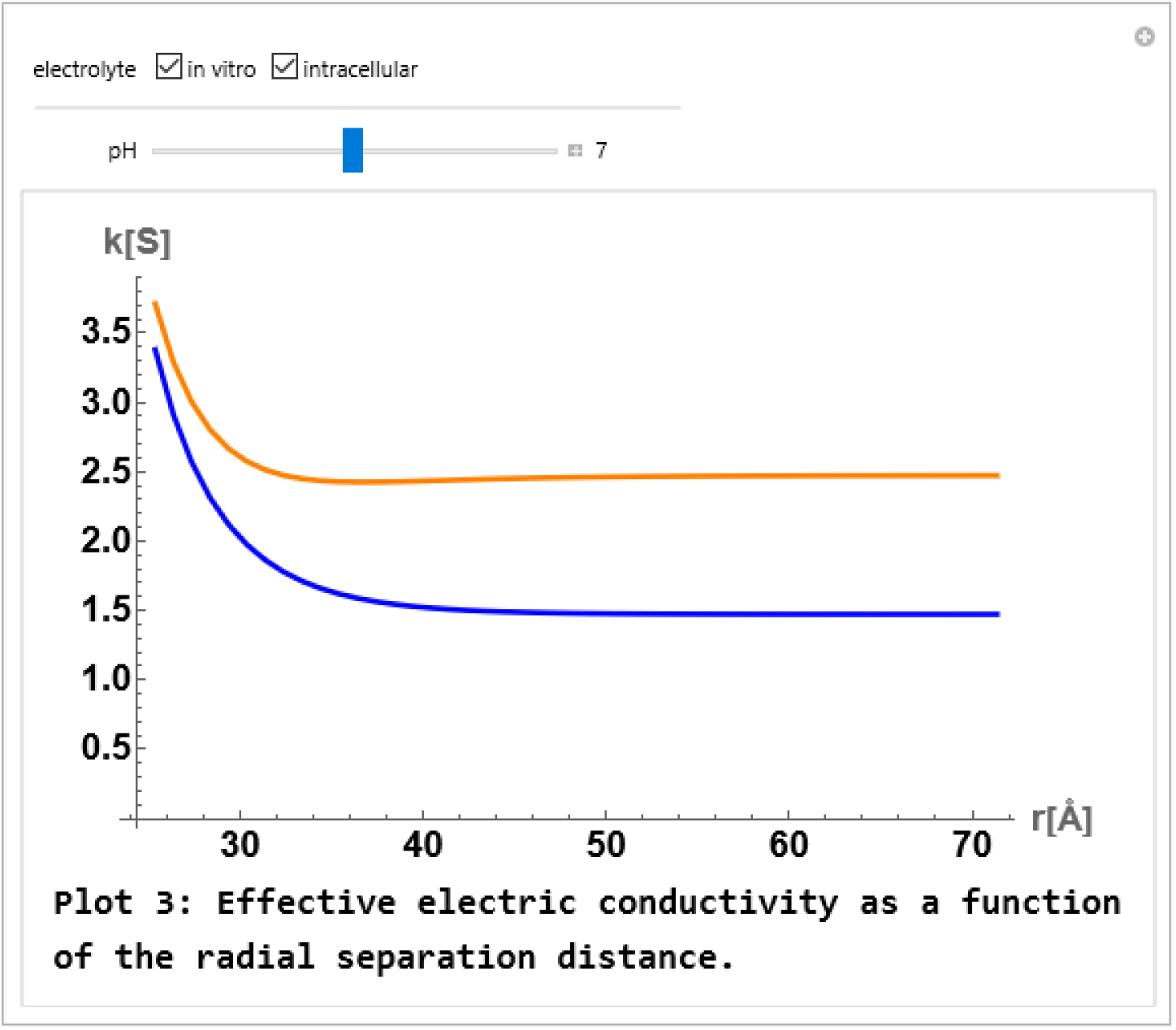
Effective filament conductivities for in-vitro(blue), and intracellular(orange) for pH 7.

#### B.2 Radial electric potential

The absolute value of the linearized Poisson-Boltzmann radial electric potential as a function of the distance is displayed in figure27 for both electrolyte conditions. The subfigure 27a reveals a higher electric potential magnitude in the intracellular condition (orange curve) when compared to in vitro condition (blue curve). The slider was used to increase the pH value showing an increasing relationship with the magnitude of the electric potential (subfigure 27b). This is an expected result, because the increase of the pH removes hydrogen ions from amino acid residues and exposes more negative charges on the protein.

**Figure 27:**
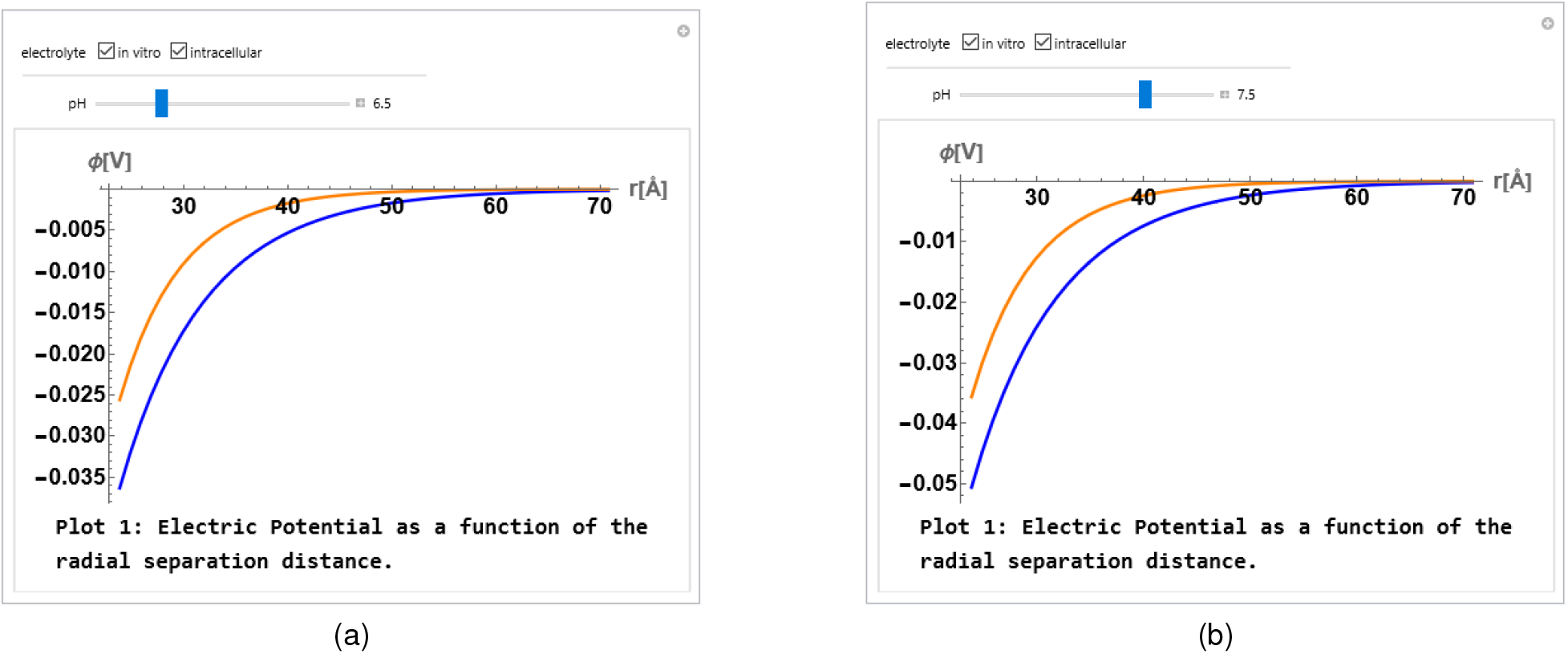
Absolute value of electrical potential as function of distance from the filament axes. Here plots for pH 6.5 (a) and pH 7.5 (b) are shown by using the slider. The orange and bluelines correspond to the intracellular and in vitro conditions, respectively.

**Figure 28:**
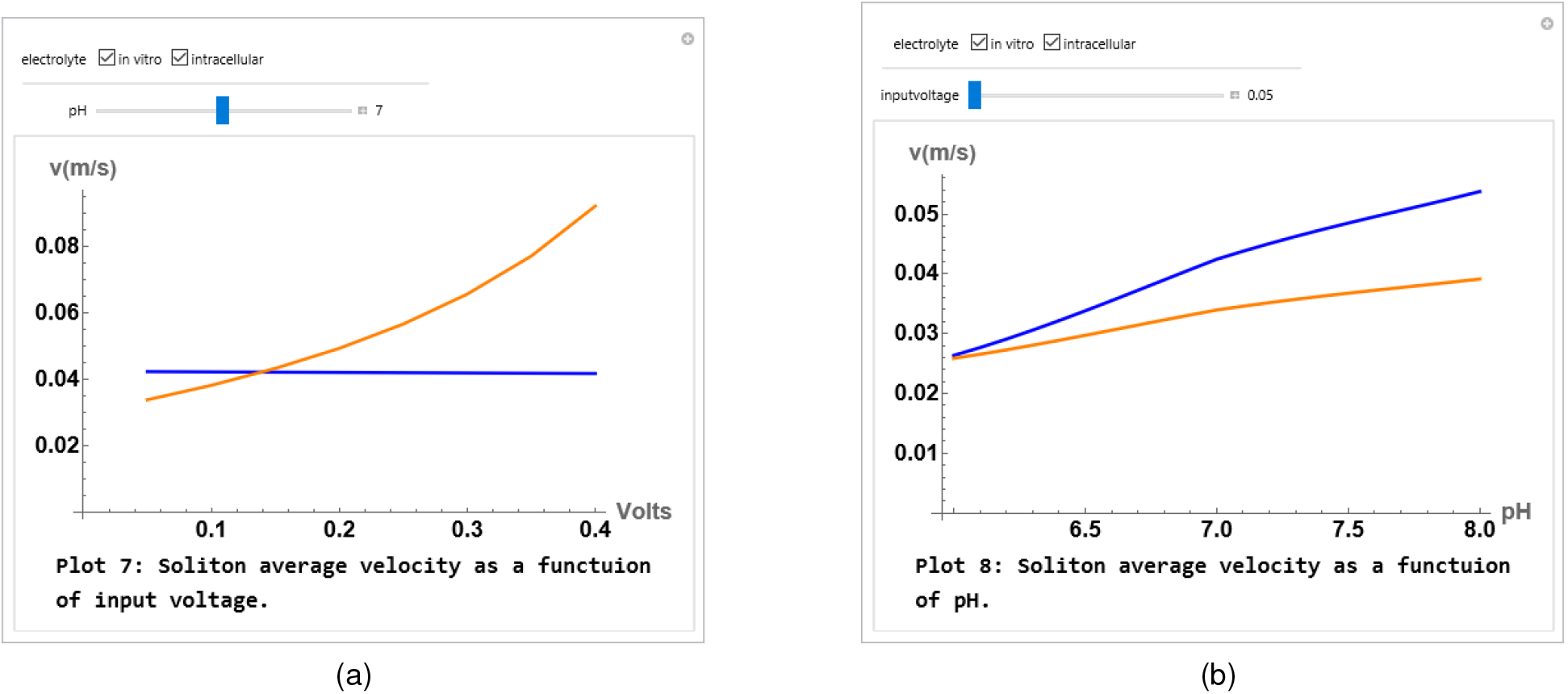
Average velocities as functions of input voltage for pH 7 (a) and as function of pH for input voltage 0.05V (b). The orangeand blue lines correspond to the intracellular and in-vitro conditions, respectively.

#### B.3 B.3 Velocity profiles

The time average soliton velocities are shown as functions of the voltage input and pH for both electrolyte conditions in subfigures 28a and 28b, respectively. With increasing input voltage, the average velocity remains almost constant for the in vitro condition. Whereas, the average velocity is enhanced with increasing input voltage for the intracellular condition. Additionally, the average velocity increases with pH for the in vitro condition at a higher rate relative to the intracellular condition. A decay in the velocity over time is demonstrated in figure 29, and concludes that initially solitons travel faster under intracellular conditions when compared to intracellular conditions, however they decelarate faster and eventually become slower than those under in-vitro conditions.

**Figure 29:**
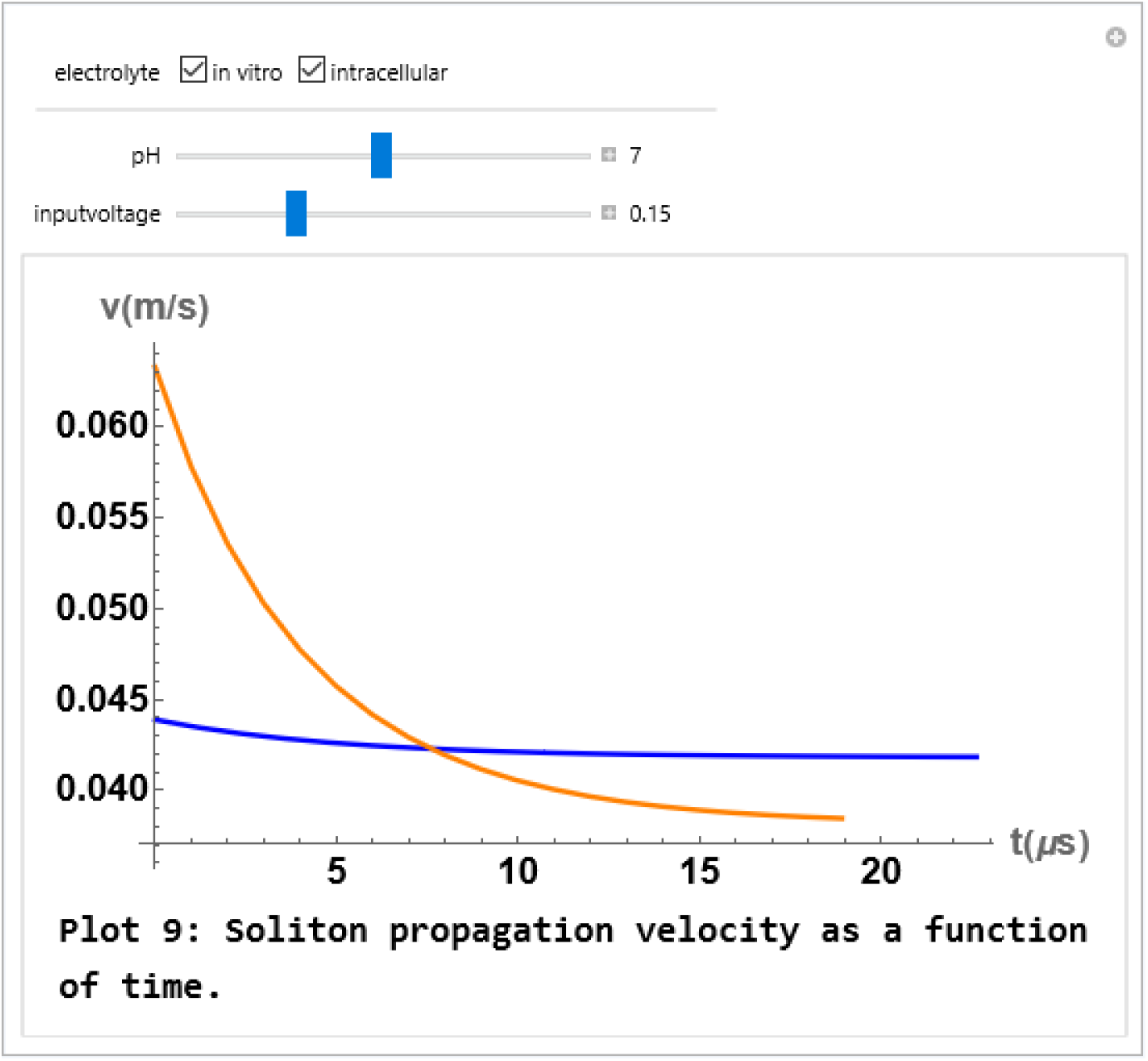
Velocity as a function of time. The orange solid and blue dashed lines correspond to the intracellular and in-vitro conditions, respectively.

#### B.4 Amplitude attenuation profiles

The first amplitude plot (Figure 30) demonstrates the soliton attenuation while allowing for the input voltage and pH to be adjusted using the slider. The peak of a decreasing soliton profile over time can be analyzed in figure 30. Here the pH of the solution has been adjusted between 6 and 8. For intracellular conditions, a lower pH value results in a faster decay of the soliton due to a weaker electrostatic screening of the EDL (Figure 30a). The result of an increased pH is shown in subfigure 30b, where a soliton peak travels further for intracellular conditions.

**Figure 30:**
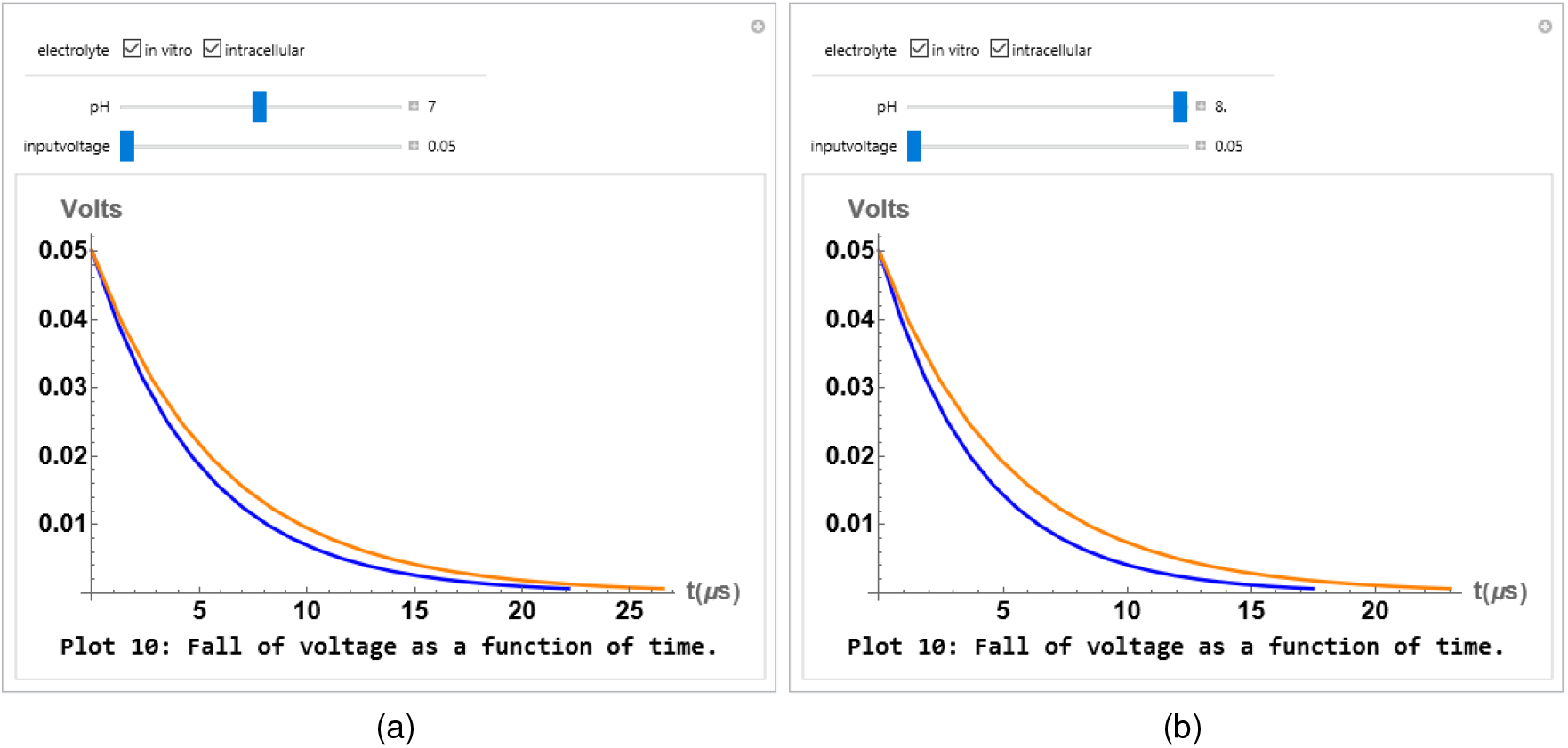
Soliton peak attenuation as a function of time at a voltage input of 0.05V and for pH 7 (a) and pH 8 (b) for in-vitro(blue) and intracellular(orange) conditions.

**Figure 31:**
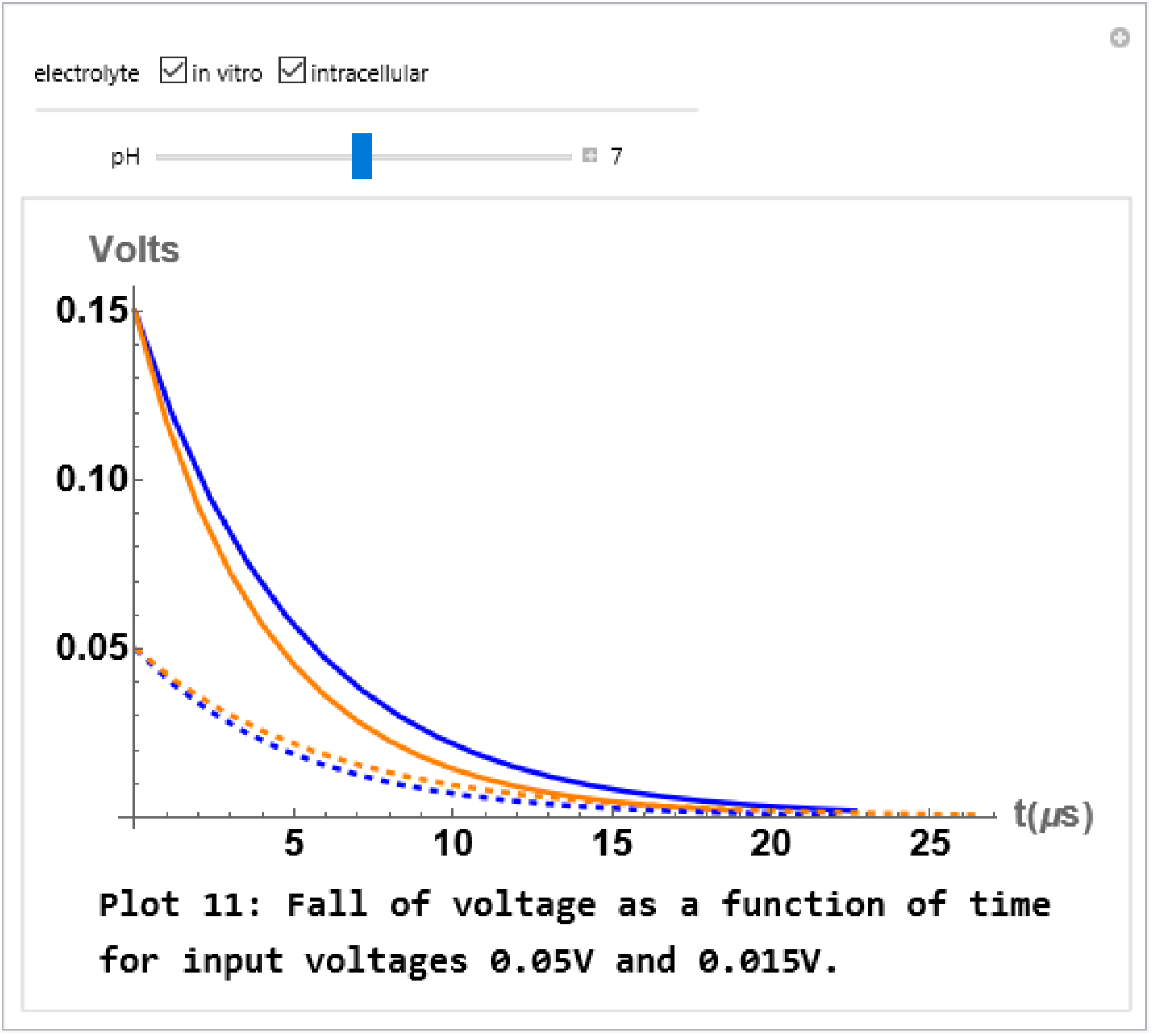
Attenuation of input voltages 0.15 V (solid lines) and 0.05 V (dashed lines) for in-vitro (blue color) and intracellular (orange color) conditions and pH 7.

The second amplitude attenuation plot (Figure 31) shows the decay of two different soliton peaks with voltage inputs *V*_0_ = 0.05*V* (dashed curves) and *V*_0_ = 0.15*V* (solid curves) for in-vitro (blue color) and intracellular (orange color) conditions and pH 7. It can be seen that larger input voltages generate faster amplitude decay rates.

#### B.5 Soliton animation profiles

By clicking the button (▶), the animations in figures 32, 33, and 34 show how a soliton propagates down the filament length. Figure 32 includes two snapshots of in vitro soliton animations at pH 7. On the other hand, figure 33 includes snapshots of the soliton propagation in intracellular conditions which initially show a faster soliton at higher voltage input but decelerate faster to eventually become slower than lower voltage input. In figure 34, we show the impact of pH on the soliton profiles in in vitro (a) and intracellular (b) conditions. We noticed that pH does not affect the soliton propagation in in vitro conditions. Whereas, a slightly faster soliton is produced at low pH in intracellular conditions. It can be seen that larger voltage inputs generate higher amplitudes, which is in agreement with experimental studies in invitro conditions [6].

**Figure 32:**
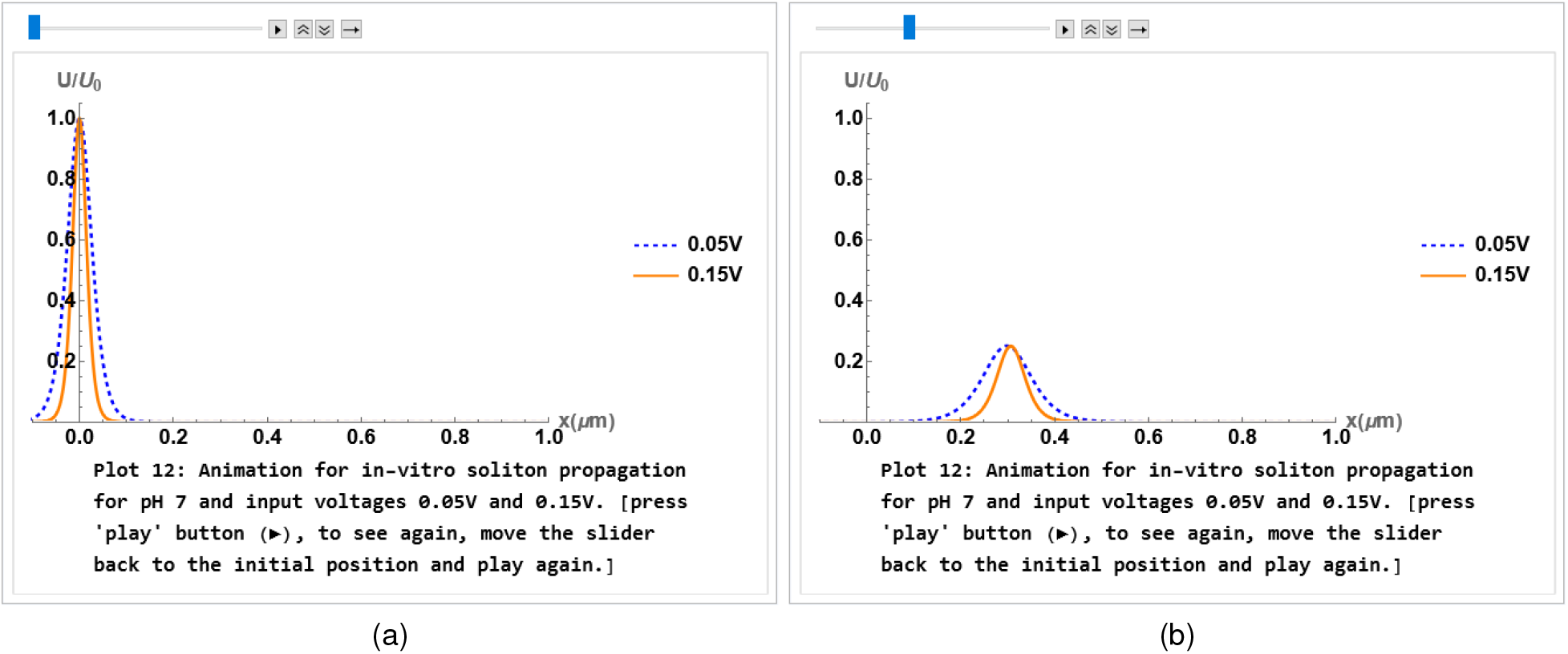
Snapshots of soliton propagation animation in in-vitro condition at the beginning (a) and at an intermediate (b) distance along the filament. Both input pulses move at same speed.

**Figure 33:**
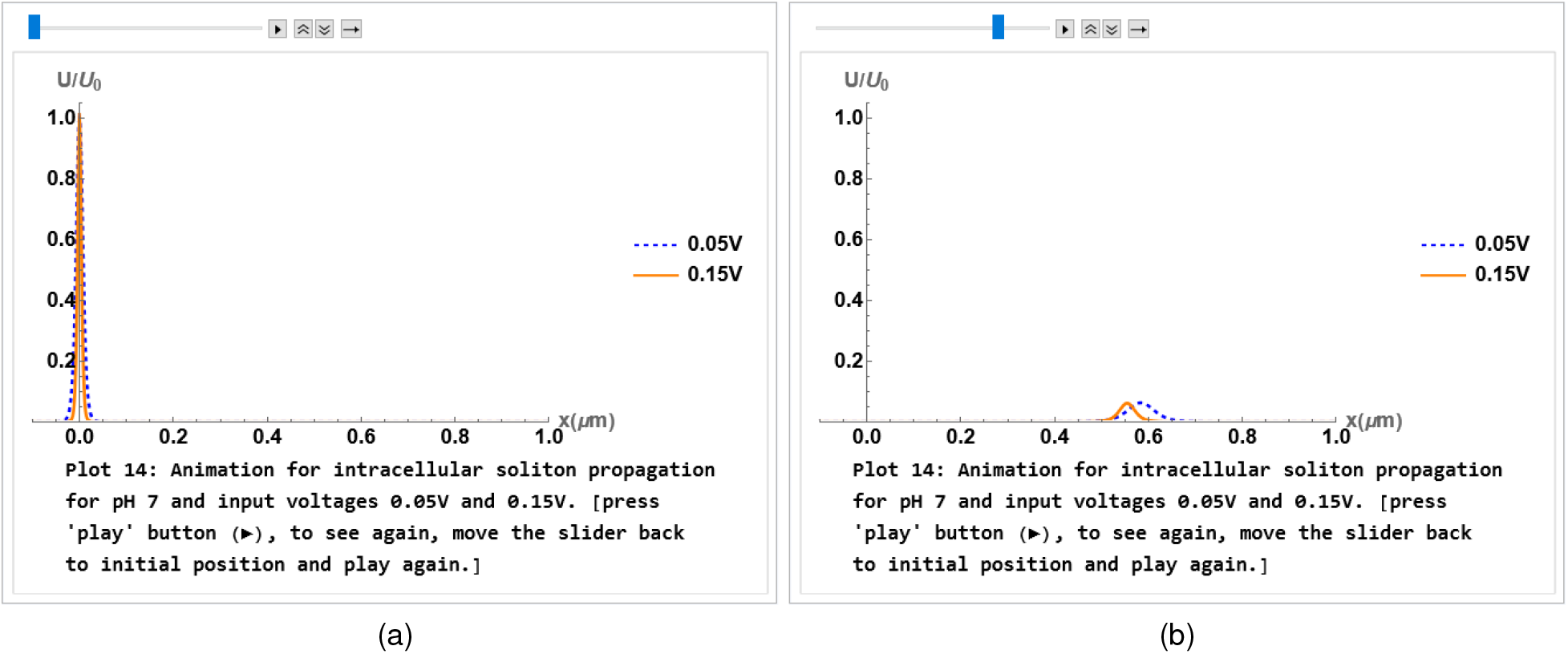
Snapshots of soliton propagation animation in intracellular condition at the beginning (a) and at an intermediate (b) distance along the filament.

**Figure 34:**
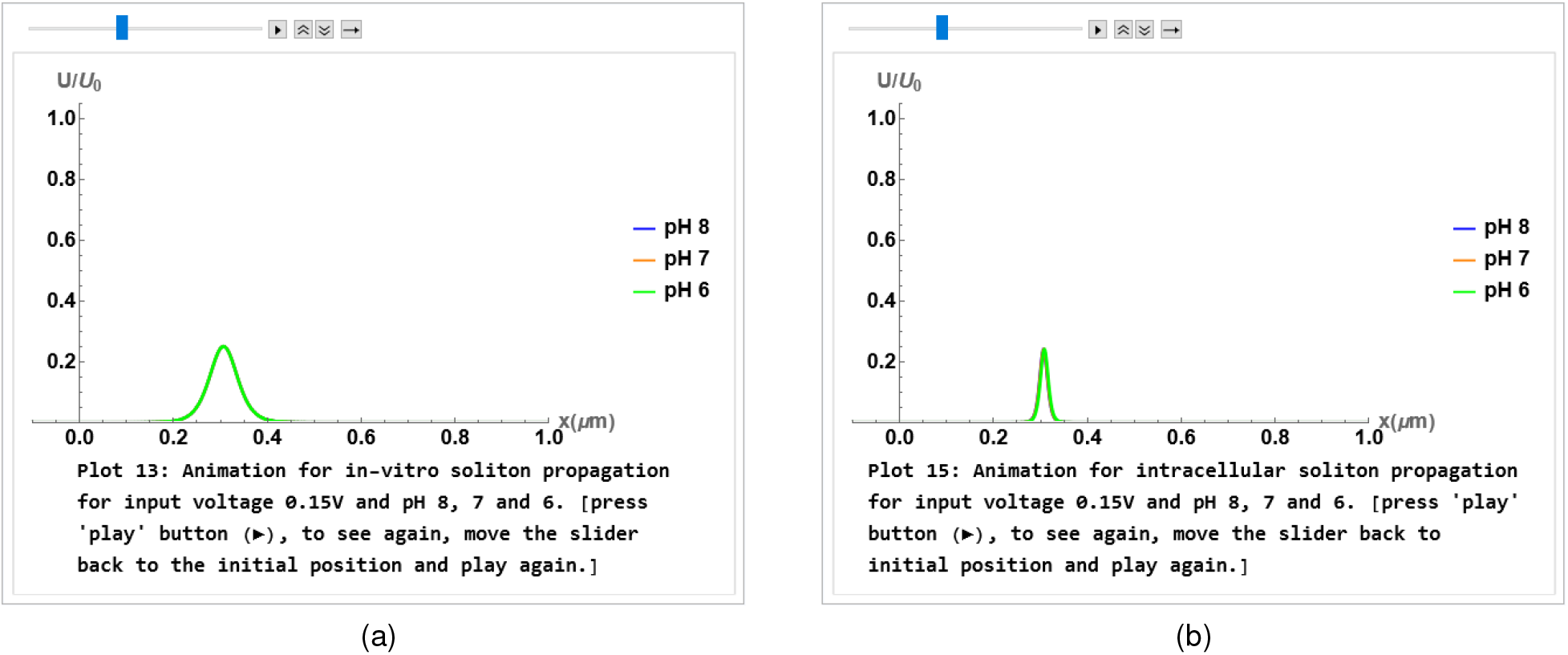
Snapshots of soliton propagation animation in in-vitro (a) and intracellular conditions at an intermediate (b) distance along the filament.

### C. Theory

In the following sections, we give a general explanation of the multi-scale approach used to produce the results for the interactive computational program. A detailed explanation on the theory can be found in a proceeding article [9].

#### C.1 The atomistic scale (Å)

The residues on the protein and pH of the solution play a large role in the formation of the EDL, which is particularly important for consideration of mutated proteins. To allow for a model that reflects changes at the atomistic scale, we performed titration curves on the most recent actin models in the protein data bank, and used a zeta potential comparison to determine the most accurate description based on the polyelectrolyte nature of actin. The Cong model (3B5U) [21] was then uploaded into the pdb2pqr webserver to determine the pH dependent amino acids exposed on the protein. The volume of the filament was determined by using the “3v: voss volume voxelator” webserver, which was then used to calculate the radius (*R* = 23.83Å), linear charge density (*λ*), and the surface charge density (*σ*).

##### C.1.1 Mutations, isoforms, and nucleotide states

Mutations can be found in all of the six human actin genes, and most often result in a substitution of an amino acid due to the change in a single base pair (missense mutation) [33]. The removal of a pH dependent residue has an effect even at the monomer scale, which is compounded in filament form by the number of added monomers. The replacement of a residue with another pH dependent amino acid will change the capacitance and both resistances for a given pH value.

#### C.2 The monomeric scale (nm)

The intrinsic properties of the actin monomer relate to the charge build up near the surface, and the ionic current next to the polymer. This can be described by an RLC circuit model, where *R* represents the resistor, *C* is the capacitor, and *L* is the inductor. The inductance has been shown to be negligible, and therefore, is not considered in the multi-scale approach [9].

##### C.2.1 Ion profiles in the electrical double layer

A detailed description of the ionic behavior in the electrical double layer is described using non-linear Boltzmann distribution of ionic concentrations.

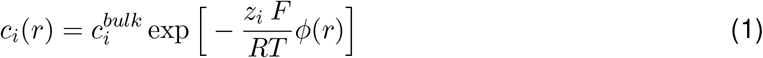

This equation describes the concentration of an ion between the surface of the actin and the bulk layer by using the ion concentration in the bulk 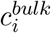, ion valance *z_i_*, Faraday’s constant F, and a scaling by the energy *RT*, where *R* is the gas constant and *T* the temperature.

Using the surface charge density (*σ*), Debye length 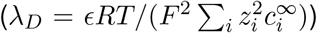, and dielectric permittivity (*ϵ* = *ϵ_o_ϵ_r_*), along with the Bessel function of the second kind gives the potential energy

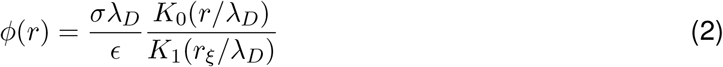

The parameter *r_ξ_* is the position of the zeta potential or slipping plane, and *r* is a distance from the surface larger than the actin radius.

The ionic current is derived from the ionic flow equation

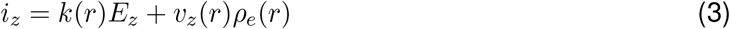

where the electric field in the longitudinal (axial) direction *E_z_* is a result of the voltage input divided by the monomer length *V*_0_/*ℓ,* and *k*(*r*) is the effective electric conductivity given by

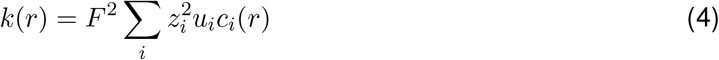

The axial ion velocity profile is given by

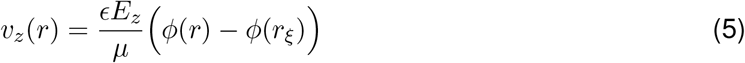

where *μ* is the viscosity described in section 2.2.1.

The ionic charge density distribution is represented by

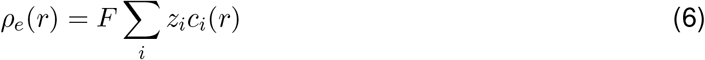

Integration over the ionic flow equation gives ion current density profile in the longitudinal direction when using *r_ξ_* and *l_B_* + *r_ξ_* as the limits of integration with *r* as the integration variable 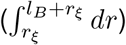, where *l_B_* = *e*^2^/(4*πϵ_o_k_B_T*) is the Bjerrum length. Therefore, we have

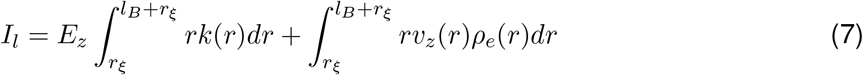

On the other hand, Integration over the ionic flow equation in the transversal (radial) direction gives the radial current density from the solution of

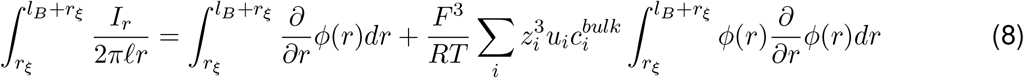

The parameter *ℓ* is the diameter of an actin monomer and *u_i_* is the ion mobility of species *i*.

The numerical solution to these integrals, which contains the non-linear predictions from the Boltzmann statistics, were performed using mathematica software [11].

##### C.2.2 Resistance

From the solution of the ion current density profiles *I_l_* and *I_r_*, Ohm’s law can be used along with the voltage drop Δ*V* =*E_z_/ℓ* to get the longitudinal and transversal resistances

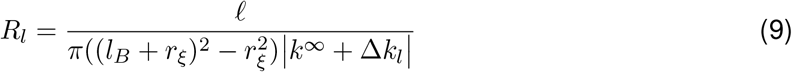

and

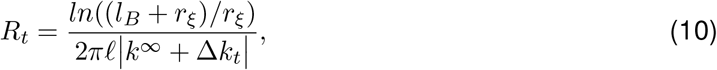

respectively.

##### C.2.3 Capacitance

Capacitance arises from the build up of counterion charge near its negatively charged surface. Therefore, the short range forces of water crowding, ion size asymmetry, and ion-ion repulsion cannot be ignored. To include these, CSDFT calculations were used for an accurate consideration of the surface electric potential *ψ_o_* = *ϕ*(*R*). This process was explained in the supplement of a previous work [9], and can easily be done by using JACFC web application [24, 25, 26]. .

The differential capacitance representing a monomer is found to be

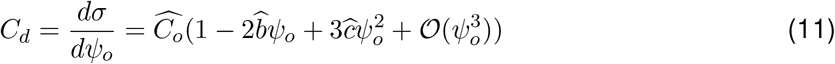

For the total charge we integrate over the voltage drop and get

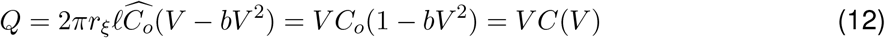

where *b* characterizes the non-linear behavior of the capacitance.

#### C.3 The filament scale (μm)

The flow of ions *I_l_*(*x, t*) = *V*(*x, t*)/*Z* along a polymerized actin filament comes from the transfer of ions from one unit cell (actin monomer) to the next. This is accomplished by using Kirchhoff’s laws on the discrete transmission line composed of N unit cells, where N represents the number of monomers in the filament. We then use a continuum approximation and a Taylor series expansion to get the perturbed Korteweg-de Vries (pKdV) differential equation

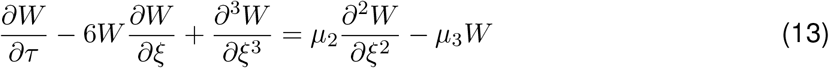

where 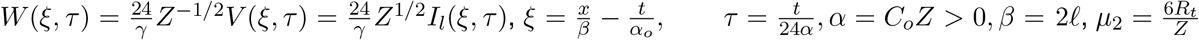 and 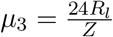.

The approximate analytic solution for the soliton is

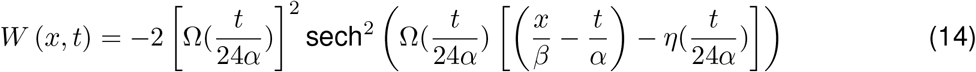

where

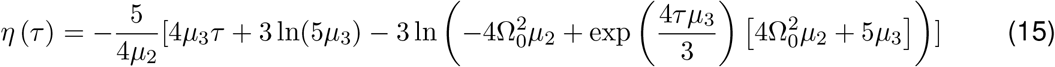

and

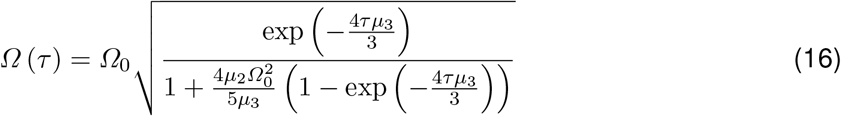

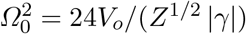 with 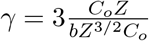

The soliton propagation velocity is given by

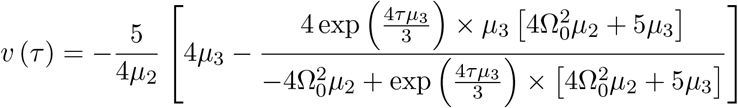

The expression for the impedance is 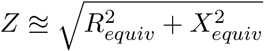, where 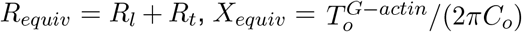 and 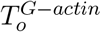 characterizes the time scale of the circuit unit.

Other important considerations are the maximum travel distance, average velocity and vanishing time represented by *x_max_, v_avg_*, and t_max_, respectively. The maximum travel distance is given by

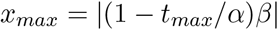

and the average velocity is

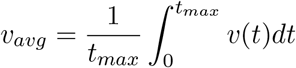

The vanishing time is the solution of the implicit equation 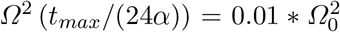, which represents the time when the soliton amplitude attenuates to 1% of its initial amplitude.

## Acknowledgements

This work was supported by the National Institutes of Health [grant number 1SC1GM127187].

## References

[1] Maria Giuseppa Leggio, Laura Mandolesi, Francesca Federico, Francesca Spirito, Benedetta Ricci, Francesca Gelfo, and Laura Petrosini. Environmental enrichment promotes improved spatial abilities and enhanced dendritic growth in the rat. Behavioural Brain Research, 163(1):78–90, 2005.

[2] Peter W. Hickmott and Iryna M. Ethell. Dendritic plasticity in the adult neocortex. The Neuroscientist, 12(1):16–28, 2006.

[3] Sara Geraldo and Phillip R. Gordon-Weeks.Cytoskeletal dynamics in growth-cone steering. Journal of Cell Science, 122(20):3595–3604, 10 2009.

[4] Thomas D. Pollard and John A. Cooper. Actin, a central player in cell shape and movement. Science, 326(5957):1208–1212, 2009.

[5] Roberto Dominguez and Kenneth C. Holmes. Actin structure and function. Annual Review of Biophysics, 40(1):169–186, 2011.

[6] H.F. Cantiello E.C. Lin. A novel method to study the electrodynamic behavior of actin filaments. evidence for cable-like properties of actin. Biophysical Journal, 65(4):1284–1289, 1993.

[7] B. R. Frieden and R. A. Gatenby. Signal transmission through elements of the cytoskeleton form an optimized information network in eukaryotic cells. Scientific Reports, 9(1):6110, April 2019.

[8] Pengyu Ren, Jaehun Chun, Dennis G. Thomas, Michael J. Schnieders, Marcelo Marucho, Jiajing Zhang, and Nathan A. Baker. Biomolecular electrostatics and solvation: a computational perspective. Quarterly Reviews of Biophysics, 45(4):427–491, November 2012. Publisher: Cambridge University Press.

[9] Christian Hunley, Diego Uribe, and Marcelo Marucho. A multi-scale approach to describe electrical impulses propagating along actin filaments in both intracellular and in vitro conditions. RSC Adv., 8:12017–12028, 2018.

[10] P. G. Drazin and R. S. Johnson. Solitons: An Introduction. Cambridge Texts in Applied Mathematics. Cambridge University Press, 2 edition, 1989.

[11] Wolfram Research, Inc. Mathematica, Version 12.3.1. Champaign, IL, 2021.

[12] Kohki Okabe, Noriko Inada, Chie Gota, Yoshie Harada, Takashi Funatsu, and Seiichi Uchiyama. Intracellular temperature mapping with a fluorescent polymeric thermometer and fluorescence lifetime imaging microscopy. Nature Communications, 3(1):705, 2012.

[13] Ryuichi Tanimoto, Takumi Hiraiwa, Yuichiro Nakai, Yutaka Shindo, Kotaro Oka, Noriko Hiroi, and Akira Funahashi. Detection of temperature difference in neuronal cells. Scientific Reports, 6(1):22071, 2016.

[14] Dominique Chretien, Paule Benit, Hyung-Ho Ha, Susanne Keipert, Riyad El-Khoury, Young-Tae Chang, Martin Jastroch, Howard Jacobs, Pierre Rustin, and Malgorzata Rak. Mitochondria are physiologically maintained at close to 50 c. PLOS Biology, 16(1):0, 2018.

[15] D. S. Viswanath, editor. Viscosity of liquids: theory, estimation, experiment, and data. Springer, Dordrecht, 2007. OCLC: ocm77482252.

[16] Ove Sten-Knudsen. Biological membranes: Theory of trans-port, potentials and electrical impulses. Cambridge Univ. Press., Cambridge, United Kingdom, 2002.

[17] Allan N. Soriano, Arjay M. Agapito, Loui John Lee I. Lagumbay, Alvin R. Caparanga, and Meng-Hui Li. Diffusion coefficients of aqueous ionic liquid solutions at infinite dilution determined from electrolytic conductivity measurements. Journal of the Taiwan Institute ofChemical Engineers, 42(2):258–264, 2011.

[18] Wolfram research, interpolation, wolfram language function. https://reference.wolfram.com/language/ref/Interpolation.html, 1991. updated 2008.

[19] P. Vanysek. Ionic conductivity and diffusion at infinite dilution. CRC Hand of Chemistry and Physics, pages 5–92, 1993.

[20] D G Miller. Estimation of tracer diffusion coefficients of ions in aqueous solution. 9 1982.

[21] Yao Cong, Maya Topf, Andrej Sali, Paul Matsudaira, Matthew Dougherty, Wah Chiu, and Michael F. Schmid. Crystallographic conformers of actin in a biologically active bundle of filaments. Journal of Molecular Biology, 375(2):331–336, 2008.

[22] Laurent Schwartz, Sabine Peres, Mario Jolicoeur, and Jorgelindo da Veiga Moreira. Cancer and alzheimers disease intracellular ph scales the metabolic disorders. Biogerontology, 21(6):683–694, 2020.

[23] Salvador Harguindey, Daniel Stanciu, JesÃºs Devesa, Khalid Alfarouk, Rosa Angela Cardone, Julian David Polo Orozco, Pablo Devesa, Cyril Rauch, Gorka Orive, Eduardo Anitua, SÃ©bastien Roger, and Stephan J. Reshkin. Cellular acidification as a new approach to cancer treatment and to the understanding and therapeutics of neurodegenerative diseases. Seminars in Cancer Biology, 43:157–179, 2017. The new pH-centric Anticancer Paradigm in Oncology and Medicine.

[24] M. marucho, university of texas at san antonio. http://neuronanobiophysics.utsa.edu. online; accessed 19-Jul-2021.

[25] Marcelo Marucho. A java application to characterize biomolecules and nanomaterials in electrolyte aqueous solutions. Computer Physics Communications, 242:104–119, 2019.

[26] Marcelo Marucho. Java application for cytoskeleton filament characterization (JACFC). Software Impacts, 8:100072, May 2021.

[27] Francine Parker, Thomas Baboolal, and Michelle Peckham. Actin mutations and their role in disease. International Journal of Molecular Sciences, 21(9):3371, 2020.

[28] Dmitri S. Kudryashov and Emil Reisler. Atp and adp actin states. Biopolymers, 99(4):245–256, 2013.

[29] Benjamin J. Perrin and James M. Ervasti. The actin gene family: Function follows isoform. Cytoskeleton, 67(10):630–634, 2010.

[30] Kristen Nowark, Gianina Ravenscroft, and Nigel Laing. muscle alpha-actin diseases (actinopathies) pathology and mechanisms. Acta Neuropathological, 125(1):19–32, 2013.

[31] Nancy Woolf, Avner Priel, and Jack Tuszynski. Nanoneuroscience: Structural and Functional Roles of the Neuronal Cytoskeleton in Health and Disease. Springer, Verlag New York, 2009.

[32] H.F. Cantiello, C. Patenaude, and K. Zaner. Osmotically induced electrical signals from actin filaments. Biophysical Journal, 59(6):1284–1289, 1991.

[33] Henderson CA, Gomez CG, Novak SM, Mi-Mi L, and Gregorio CC. Overview of the muscle cytoskeleton. Comprehensive Physiology, 7(3):891–944, 2017.

